# What Large Language Models Know About Plant Molecular Biology

**DOI:** 10.1101/2025.08.31.672925

**Authors:** Manuel Fernandez Burda, Lucia Ferrero, Nicolás Gaggion, Camille Fonouni-Farde, The MoBiPlant Consortium, Martín Crespi, Federico Ariel, Enzo Ferrante

## Abstract

Large language models (LLMs) are rapidly permeating scientific research, yet their capabilities in plant molecular biology remain largely uncharacterized. Here, we present MOBIPLANT, the first comprehensive benchmark for evaluating LLMs in this domain, developed by a consortium of 112 plant scientists across 19 countries. MOBIPLANT comprises 565 expert-curated multiple-choice questions and 1,075 synthetically generated questions, spanning core topics from gene regulation to plant-environment interactions. We benchmarked seven leading chat-based LLMs using both automated scoring and human evaluation of open-ended answers. Models performed well on multiple-choice tasks (exceeding 75% accuracy), although most of them exhibited a consistent bias towards option A. In contrast, expert reviews exposed persistent limitations, including factual misalignment, hallucinations, and low self-awareness. Critically, we found that model performance strongly correlated with the citation frequency of source literature, suggesting that LLM knowledge inherits the visibility distribution of the underlying scientific corpus. Consequently, models tend to be more reliable on consolidated topics and less reliable on under-cited or recently emerging ones. We also benchmarked agents equipped with web-search and additional tools in more complex tasks involving DNA sequence analysis. These agents were outperformed by domain specific models in sequence classification and regression tasks, indicating an opportunity for joint agentic systems that combine both the reasoning power of LLMs and the dedicated processing of DNA models. This understanding is key to guiding both the development of next-generation models and the informed use of current tools in the everyday work of plant researchers. MOBIPLANT is publicly available online [in this link](https://huggingface.co/datasets/manufernandezbur/MoBiPlant).

## 1 Introduction

Large language models (LLMs) have demonstrated transformative potential across the research lifecycle, assisting with tasks such as automated literature surveys, data extraction, and experimental design guidance^1^. Researchers now use LLMs not only to improve writing quality, but also to interpret biological literature, propose novel research directions, assist with data analyses, and uncover functional patterns in complex biological data^2^. Despite their widespread adoption, these models still require careful expert validation to understand their capabilities and limitations, as domain-specific evaluations remain limited. To date, most large-scale evaluations focus on general-purpose or biomedical domains, leaving gaps in specialized fields like plant molecular biology^3^. Notably, LLMs have demonstrated proficiency in logical reasoning and scientific benchmarks, often surpassing previous performance records. While benchmarks like SciBench^4^, LAB-Bench^5^ and BrainBench^6^ have been developed to assess LLMs’ scientific knowledge and practical research capabilities, they primarily focus on general scientific disciplines or specific areas like biology, chemistry or neuroscience. But standardized, domain-specific evaluations remain scarce, particularly in plant molecular biology. To address this gap, we introduce the Plant Molecular Biology (MOBIPLANT) Benchmark, the first expert-curated plant-biology benchmark created by a consortium of more than 110 plant scientists residing in 19 different countries, combining manual and synthetically generated multiple-choice questions (MCQs) with rich human-evaluation of open-ended answers.

The few existing studies which survey the use of AI in research environments, underscore both excitement and caution, highlighting how researchers leverage chat assistants to accelerate literature review and experiment planning, yet worry about uncontrolled hallucinations and biorisk implications^7^. Meanwhile, frontier systems demonstrate that agentic LLMs can achieve or exceed expert performance on literature-search and contradiction-detection tasks^2,8,9^, suggesting that with the right evaluation frameworks, models could reliably support scientific discovery. In agriculture there have been approaches to leverage farmer queries through LLM-powered systems^10^. For genomics, initial studies reveal feasibility but also highlight persistent domain-specific challenges: for example, ChatGPT’s outputs exhibit “plant blindness” and limited taxonomic diversity when answering botanical questions^11^; and LLMs, while extracting ecological data more than 50 times faster than human reviewers, require rigorous quality-control to avoid quantitative errors^2^. Beyond natural language queries, plant molecular biology increasingly relies on tasks that operate directly on DNA sequences^12^, such as regulatory element detection or promoter strength prediction, where the boundary between language models and sequence models becomes relevant. Whether general-purpose LLMs, even when augmented with agentic capabilities (like web-search or coding tools), can compete with specialized sequence architectures on such tasks remains an open question.

Moreover, these models have been shown to misinterpret bioinformatics figures (such as reversing up- and down-regulated gene sets in differential expression plots) producing plausible yet incorrect biological narratives^13^. Another example is chat-based extraction of natural-product bioactivity from the literature, which yields high recall but introduces false positives, assigning activity values to compounds never tested^14^. Clinical workflows further demonstrate that, without careful oversight, LLMs can present over-confident hallucinations—such as fabricated drug interactions—or omit critical diagnostic details, reinforcing the necessity of human-in-the-loop validation^15^.

The proposed research pursues four interlocking objectives to rigorously assess LLM capabilities in plant molecular biology. First, we settled an expert Consortium composed of 112 scientists specialized in the field (Figure 1): the MoBiPlant Consortium. Second, we built an open and community-driven benchmark that unites expert-crafted, PhD-level MCQs (Expert MoBiPlant) with a synthetically expanded question set (Synthetic MoBiPlant), ensuring both the precision of domain-vetted items and the topical breadth afforded by controlled data augmentation. Third, we carried out a dual-mode evaluation: high-throughput automated scoring on the MCQs to establish a baseline accuracy, alongside in-depth human assessment of open-ended responses, examining not only correctness but also alignment with scientific consensus, potential of species bias, logical reasoning, and self-awareness (measured as the acknowledgement of limitations expressed by the model). Fourth, we evaluated the capabilities of state-of-the-art agentic LLMs on DNA sequence analysis tasks, revealing a scenario in which domain-specific DNA models decisively outperform even the most powerful frontier agents on nucleotide-level reasoning. Yet they remain unable to handle DNA tasks unseen during training, underscoring the complementary rather than substitutable roles of these two model families.

**Figure 1.**
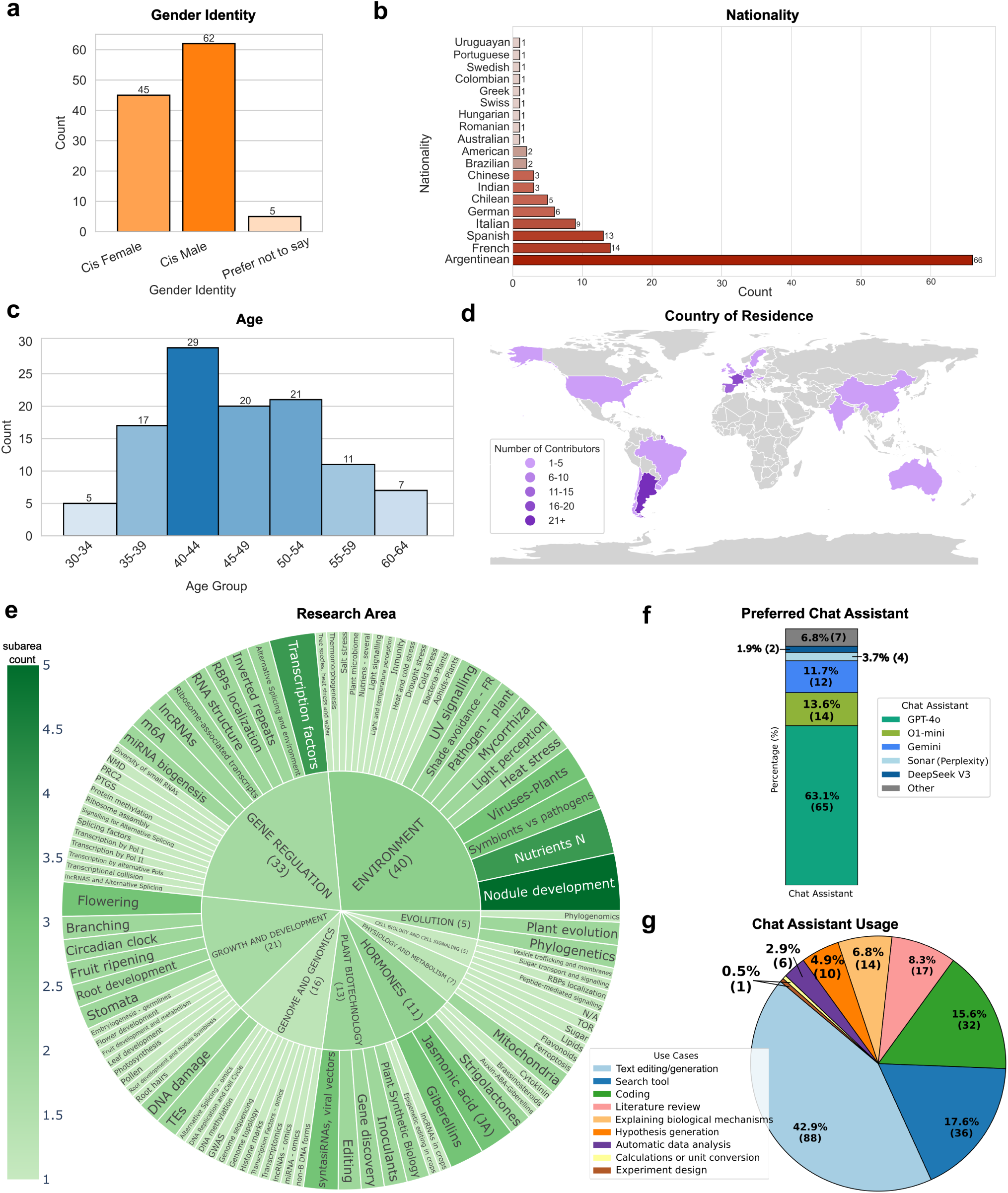
MoBiPlant Consortium Description including **a)** gender, **b)** nationality, **c)** age, **d)**, country of residence and **e)** research area. It is worth mentioning that in ‘Nationality’ and ‘Research Area’, contributors might have reported more than one answer. **f)** In addition, we queried the Consortium which LLM was their favourite to use as assistant in research contexts, **g)** and how they employed them.

We evaluated some of the most popular LLMs used as chat assistants by scientists in plant molecular biology: Gemini^16^ (1.5 Pro), ChatGPT^17,18^ (both GPT-4o and o1-mini), DeepSeek^19,20^ (both V3 and R1); a well-known open source chat assistant: LLaMA^21^ (3.1 405B); and an assistant excelling in scientific benchmarks: Claude^22^ (3.5 Sonnet). Through the MOBIPLANT framework, we uncovered model specialization patterns across domains and performance features. Finally, we offered a modular MCQ construction protocol for PhD-level question design, enabling the community to extend and adapt our benchmark to new scientific domains.

Our results highlight both the strengths and weaknesses of popular LLMs when tasked with understanding and reasoning about plant molecular biology. Despite demonstrating strong performance in structured MCQ tasks, these models still struggle with factual accuracy, hallucinations, and logical reasoning, particularly in the context of novel or less-represented scientific concepts. Notably, the fact that performance of LLMs correlates with the citation frequency of the source papers suggests that models may be better equipped to handle well-established concepts that are more likely to appear in their training data, while struggling with newer or less-represented material. Additionally, by benchmarking agentic LLMs against a dedicated DNA language model on plant tasks that involve DNA sequences, we show that general-purpose models augmented with search-tools and coding capabilities still fall short of fine-tuned sequence architectures, underscoring the complementary roles these systems are likely to play in future plant research workflows. Through our comprehensive benchmarking, we underscore the need for targeted improvements to enhance the reliability and scientific rigor of LLMs, particularly in specialized domains like plant molecular biology.

## 2 Results

### 2.1 MoBiPlant allows assessment of LLMs understanding in plant molecular biology

Aiming to understand the limitations of popular LLMs when used by plant molecular researchers in their everyday routine, we created a diverse consortium of 112 plant molecular biology researchers from around the world, referred to as the MoBiPlant Consortium (Figure 1), which was in charge of elaborating complex questions and evaluating open-ended answers (see Methods 5.1. Our evaluation methodology to assess LLMs’ understanding of plant molecular biology consisted of a two-stage experimental design: (1) question generation and (2) answer evaluation. During the question generation stage, researchers from the MoBiPlant Consortium were tasked with selecting relevant, high-quality scientific papers and creating complex, domain-specific questions based on the content of these papers (Figure 2.a). The consortium elaborated the questions, established correct answers, and then conducted a personal in-depth evaluation of LLM-generated open-ended responses. As a result, MoBiPlant is the first comprehensive, expert-driven benchmark that integrates rigorously vetted MCQs with open-ended assessments created and reviewed by domain specialists. Moreover, each question was accompanied by a correct answer and two plausible distractors, designed to form a MCQ set. These questions constituted the **Expert MoBiPlant** set.

**Figure 2.**
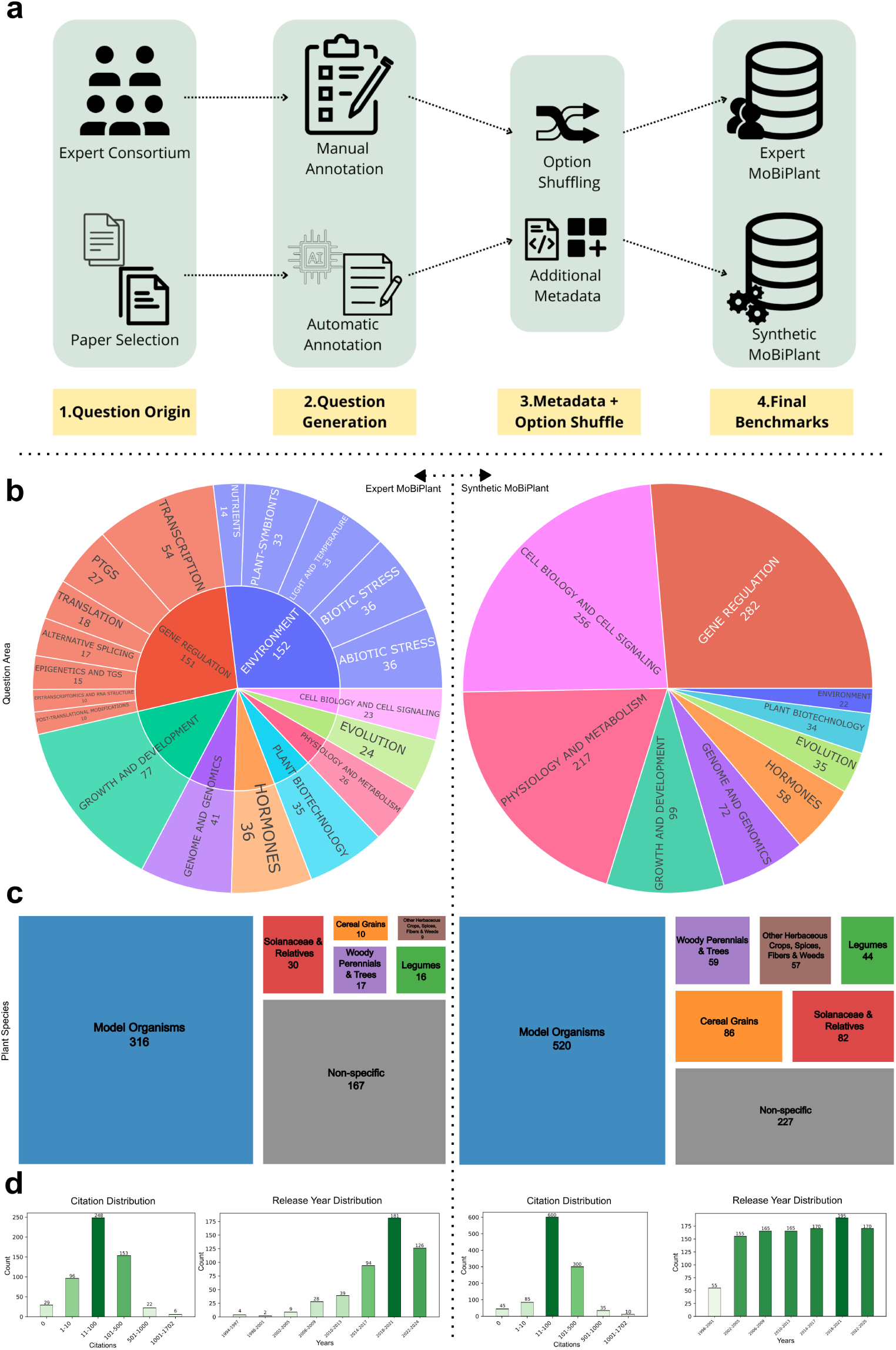
MoBiPlant Overview. **a)** MoBiPlant construction pipeline; **a.1)** The MoBiPlant Consortium was created and the source papers were selected, ensuring no overlap. **a.2)** Scientists manually created the Expert questions based on their expertise, always indicating a source paper where the answer is rooted, whereas synthetic ones were automatically generated from source papers via a frontier LLM, with question topics and plant species annotated at this stage. **a.3)** Answer options were shuffled, and metadata—such as plant species names—was standardized and cleaned. **a.4)** Both expert and synthetic benchmarks. **b)** Question area organization: the inner circle contains the main areas, while secondary areas (if applicable) are in the outer circle. **c)** Plant species distribution grouped by 6 delimiting categories. **d)** Source paper distribution according to release year and citation count.

To scale up our study and perform a more robust analysis, we constructed an additional set of 1075 synthetically generated questions derived from already published papers (Figure 2.a). More precisely, we employed a frontier LLM (see Methods 5.2) to generate synthetic MCQs based on existing scientific papers. In this process, the LLM was provided with a paper manuscript and prompted to produce questions, each with one correct answer and two incorrect alternatives. An expert group validated a subset (n=20, see details in Methods 5.2) of these questions ensuring the set generated attained to the criteria. This set of LLM-generated questions was referred to as the **Synthetic MoBiPlant** set.

In the second stage, i.e. answer evaluation, we prompted alternative popular LLMs (see Methods 5.3) to respond to both sets of questions in two distinct formats:

- **MCQ format**, where the model selected the correct answer from a predefined list of options for both Expert and Synthetic sets. Evaluation was performed automatically, by simply comparing the selected answer against the correct answers, and computing accuracy (see Methods 5.4).
- **Open-ended format**, only for the Expert MoBiPlant set, allowing the models to generate full, unconstrained responses. A detailed manual evaluation was conducted by experts from the MoBiPlant Consortium, who assessed the quality and factuality of the model-generated answers. This evaluation was based on a set of pre-defined criteria (see Methods 5.5).

### 2.2 LLMs Encode Plant Molecular Biology Knowledge

When benchmarking 7 of the most popular LLMs using the Expert MoBiPlant set in MCQ format, we see that all models achieve results over 75% accuracy (Figure 3.a). We find Claude 3.5 Sonnet to be the best performing model in our dataset with an overall score of 88.1%, highlighting its strong capabilities in agreement with previous observations on different scientific domains^23^.

**Figure 3.**
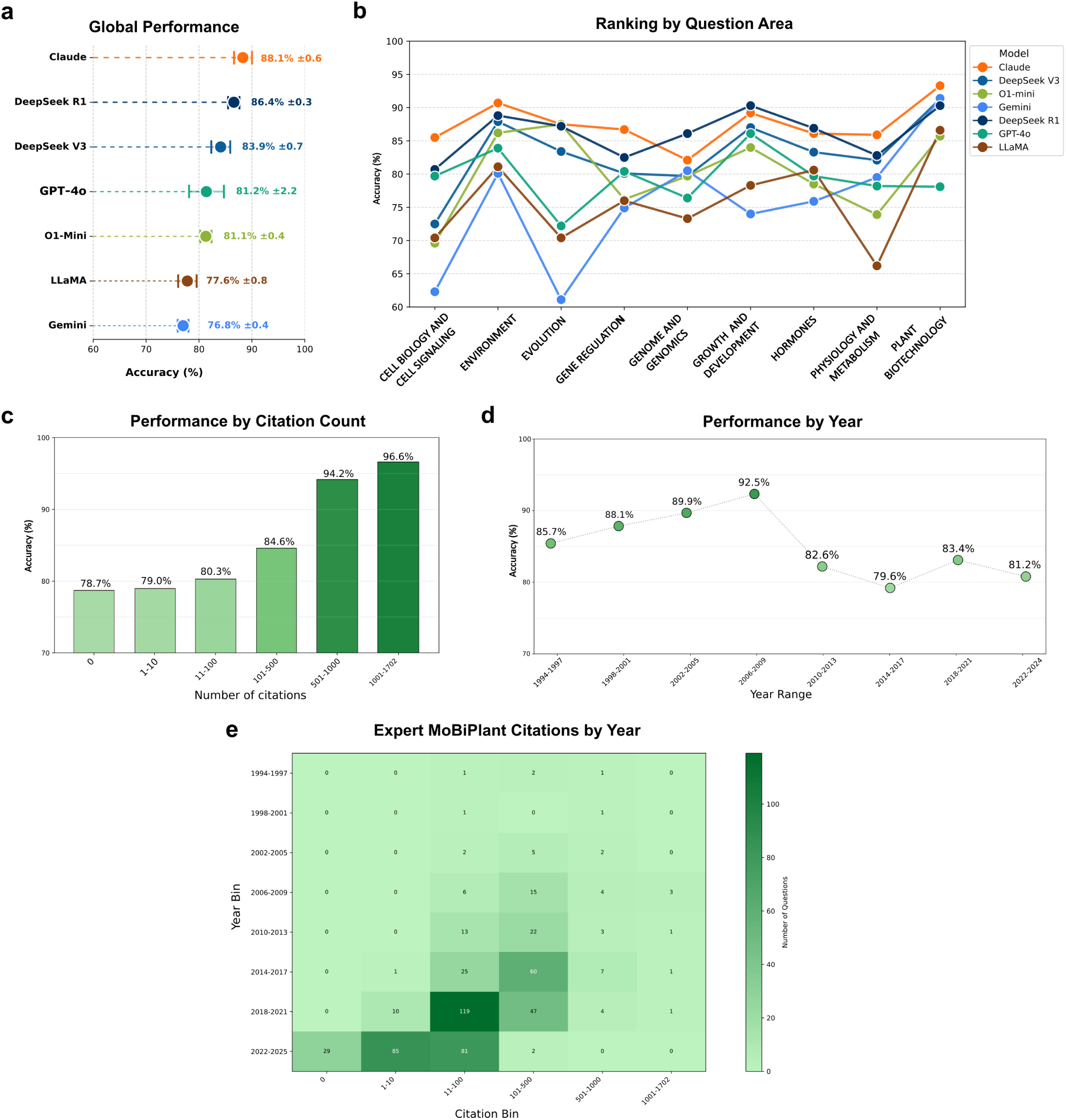
Expert MoBiPlant MCQ benchmarking results. **a)** The overall accuracy of different LLMs across Expert MoBiPlant. We report the mean value of accuracy among its standard deviation as error bars for all 3 independent repetitions of model answering, shuffling the option order on every repetition. **b)** Fluctuation of model ranking across question areas. Dots represent the model accuracy for different question areas. **c)** Performance by citation count and **d)** year, extracted from the paper that originated the question. The color intensity is proportional to the accuracy score on each plot. **e)** The represented publications broken down by citations and year of publication in Expert MoBiPlant accounted for with a heatmap, highlighting the intersected coverage between year of publication and amount of citations ranges.

The model ranking for questions coming from different research areas highlights topic specialization across LLMs (Figure 3.b). By plotting the shifts in models’ order for all the areas benchmarked in Expert MoBiPlant, it is notable that there is a correlation between the two best scoring models, i.e. Claude 3.5 Sonnet and DeepSeek R1, in the global score chart and their rank in every area, securing the first two places in most categories. This however does not hold true for every other model, where the rankings get shuffled among areas. Interestingly, this topic specialization is clearly depicted in Plant Biotechnology, where Gemini, the lowest scoring model in our benchmark, achieves a score of 91.4%, securing the second place among top performances.

To better support our observations, we assessed the performance of the same LLMs on the Synthetic MoBiPlant set of MCQs, performing identical analyses. Overall, the scores are positively shifted (Supplementary Figure 1.a), granting diverse gains in model performance; from 0.03 (Claude) to 19.7 (LLaMA) in percentage points. The overall higher accuracy suggests that this set might be easier for models to answer, highlighting the importance of expert human intervention in creating challenging questions to evaluate model understanding. This is likely because synthetic distractors are generated by tweaking the correct answer drawn from a single source paper, whereas Expert MoBiPlant distractors are crafted by specialists to reflect closely related but incorrect concepts (e.g., neighboring pathways or similar gene names). In addition, the leaderboard gets shuffled in this case with Deepseek V3, Claude and LLaMA leading. The same model specialization can be observed, as the model position significantly varies across areas (Supplementary Figure 1.b). Similar to the assessment of LLMs using the MobiPlant Expert set (Figure 3.c), referring to the performance by citation count on Synthetic MoBiPlant set results in a clear positive correlation between the amount of citations and model accuracy (Supplementary Figure 1.c).

### 2.3 Established Domain Knowledge Shapes LLM Performance

To assess how citation impact and publication year influence model performance on MoBiPlant, we analyzed questions anchored to specific source papers. For each source paper, we extracted its publication year and citation count, and stratified the data into count bins (Figure 3.c-d). Our analysis revealed a correlation between citation frequency and model accuracy: studies with citation counts in the 501–1,702 range exhibited significantly higher accuracy compared to those in the 0–500 interval. Additionally, when computing the Pearson correlation between the mean model accuracy (unbinned) and the citation count, we found a statistically significant positive correlation (r = 0.139, p-value = 1.0454 × 10^−3^). This trend likely arises because highly cited works are more frequently represented in training corpora, either through direct inclusion or repeated paraphrasing, thereby reinforcing the model’s familiarity with their content.

Interestingly, when papers were stratified by publication year, we observed a persistent trend: for articles published until 2009, the related questions were answered with higher accuracy (ranging from 87.5% to 92.5%) than those published more recently (since 2010, with the accuracy ranging from 79.6% to 83.4%). Considering the correlation between citations and accuracy (Figure 3.c), we assessed if older papers picked by the MoBiPlant members were more highly cited. However, this difference was not observed (Figure 3.e). Remarkably, the accuracy trend observed in the Synthetic MoBiPlant set (Supplementary Figure 1.d) closely mirrors that of the Expert MoBiPlant set. However, the difference in accuracy between older and more recent articles is less pronounced, ranging from 86.3% to 89% for articles published up to 2009, and from 85.1% to 86.8% for those published since 2010. Importantly, the distribution of citations across articles stratified by year of publication remains relatively uniform, with a more balanced number of articles per year compared to the Expert MoBiPlant set—thereby minimizing potential bias (Supplementary Figure 1.e). One plausible explanation for the higher accuracy of LLMs answers to questions based on older articles—despite these articles not necessarily being more highly cited— is that the knowledge they introduced has been indirectly cited and assimilated into more recent publications, becoming part of the established scientific consensus. To test this hypothesis, we segregated articles in the Synthetic MoBiPlant set into research papers and review articles. Since review articles aim to summarize, synthesize, and critically analyze the current state of knowledge on a given topic, they offer an ideal lens through which to examine the consolidation and diffusion of scientific information. By splitting the journals into two groups: review articles and research articles (papers), we calculated the accuracy on each of them (Figure 4.a). Interestingly, reviews surpass papers by 10 to 15 percentage points on each model, albeit the slight -yet significant-difference in citation distributions between reviews and papers (Figure 4.b). These findings suggest that review articles tend to reflect more consolidated knowledge which is better assimilated by LLMs. Accordingly, review publications often synthesize well-established conceptual frameworks that form part of the shared understanding in plant molecular biology, resulting in clearer and more universally accepted ideas. Taken together, our analyses of citation trends and the integration of key concepts in review articles underscore the role of canonical knowledge in shaping LLM performance in the domain of plant molecular biology.

**Figure 4.**
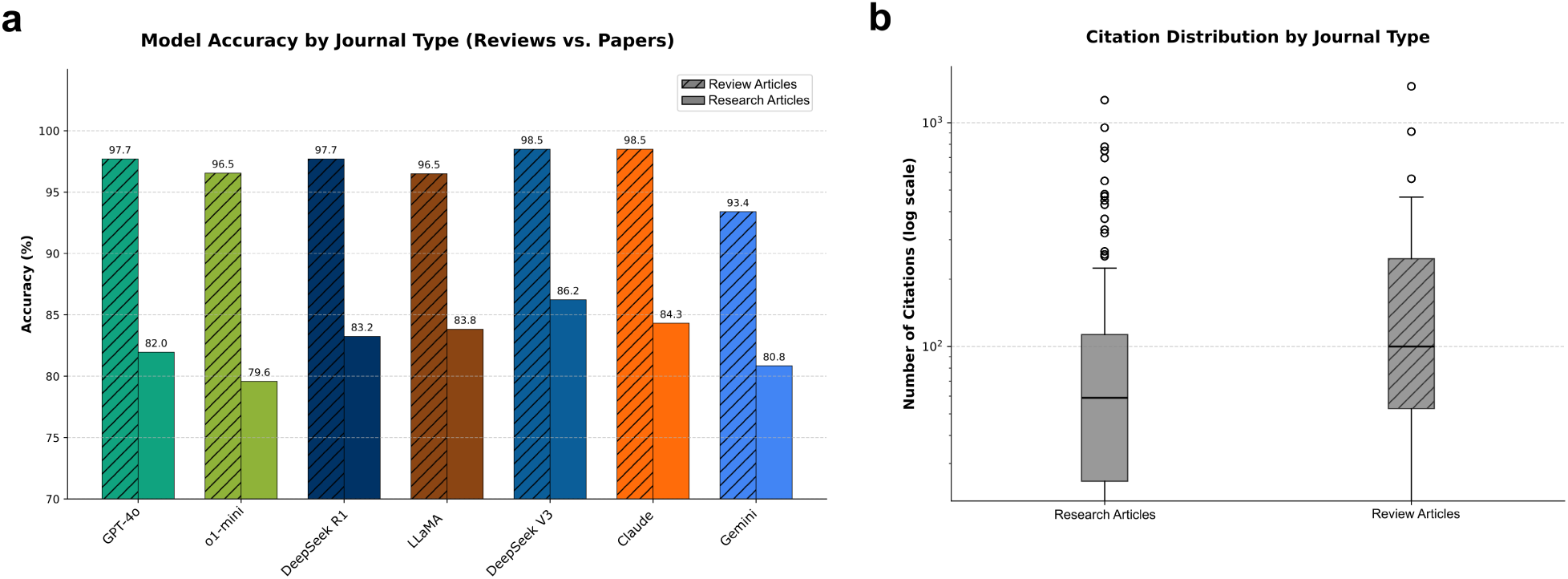
Performance of LLMs based on the type of journal used as source of information. Source journals are grouped by their type of publication, either review articles or research articles. **a)** Accuracy by model on each journal, where the dashed bar represents the mean accuracy over review journals and the solid bar represents the mean accuracy over the paper journals. **b)** The citation distribution among these two groups, represented in boxplots. Y-axis: citations on logarithmic scale. We further verified if there was a citation bias on these groups computing the Mann-Whitney U-test with a p-value of 4.536 × 10^−13^, suggesting a statistically significant difference in citation distributions between review and paper journals.

### 2.4 Answer Order Influences LLM Performance On MCQs

As explained in the Methods section, answer options in the MQCs were randomly shuffled into three groups to minimize positional bias. While approximately 33% of the correct answers were assigned to option A in each group, five out of seven from the evaluated models selected more than 37% of the time the option A, with LLaMA reaching over 45% (Figure 5.a). Remarkably, O1-mini and Claude 3.5 Sonnet did not exhibit any bias towards the option A. This observation suggests that when uncertain, the rest of the LLMs tend to default to option A, potentially inflating their performance metrics or obscuring their true accuracy. Considering that LLMs previously showed bias towards the order of answers in MCQs^24,25^, we conducted a parallel experiment on models’ election distribution (A,B or C) to explore potential option-bias risks. We generated 3 distinct versions of Expert MoBiPlant, reordering the options so that the correct option was fixed in a particular choice (the first having every correct option in A, the second in B and the third in C). Then, we evaluated one of the best models and one of the worst models in Expert MoBiPlant (Figure 5.b, Supplementary Table 2). Overall, we see that both models have a bias towards option A (although DeepSeek R1 seems to be more robust to these shuffles while still showing flaws). We extend on this by denoting: 1) accuracy tops up when the correct option is set on A (even surpassing the vanilla Expert MoBiPlant performance) and that when the correct option is not on A, accuracy consistently drops (Supplementary Table 2); 2) model’s choice tends to be A even when it is not the correct option; on the second (all corrects are B) and third (all corrects are C) versions Gemini chooses three times more the option A than its other wrong counterpart (while DeepSeek shows a less pronounced tendency). This settles the ground for posterior discussion about complementing MCQ evaluation with robust human reviewing.

**Figure 5.**
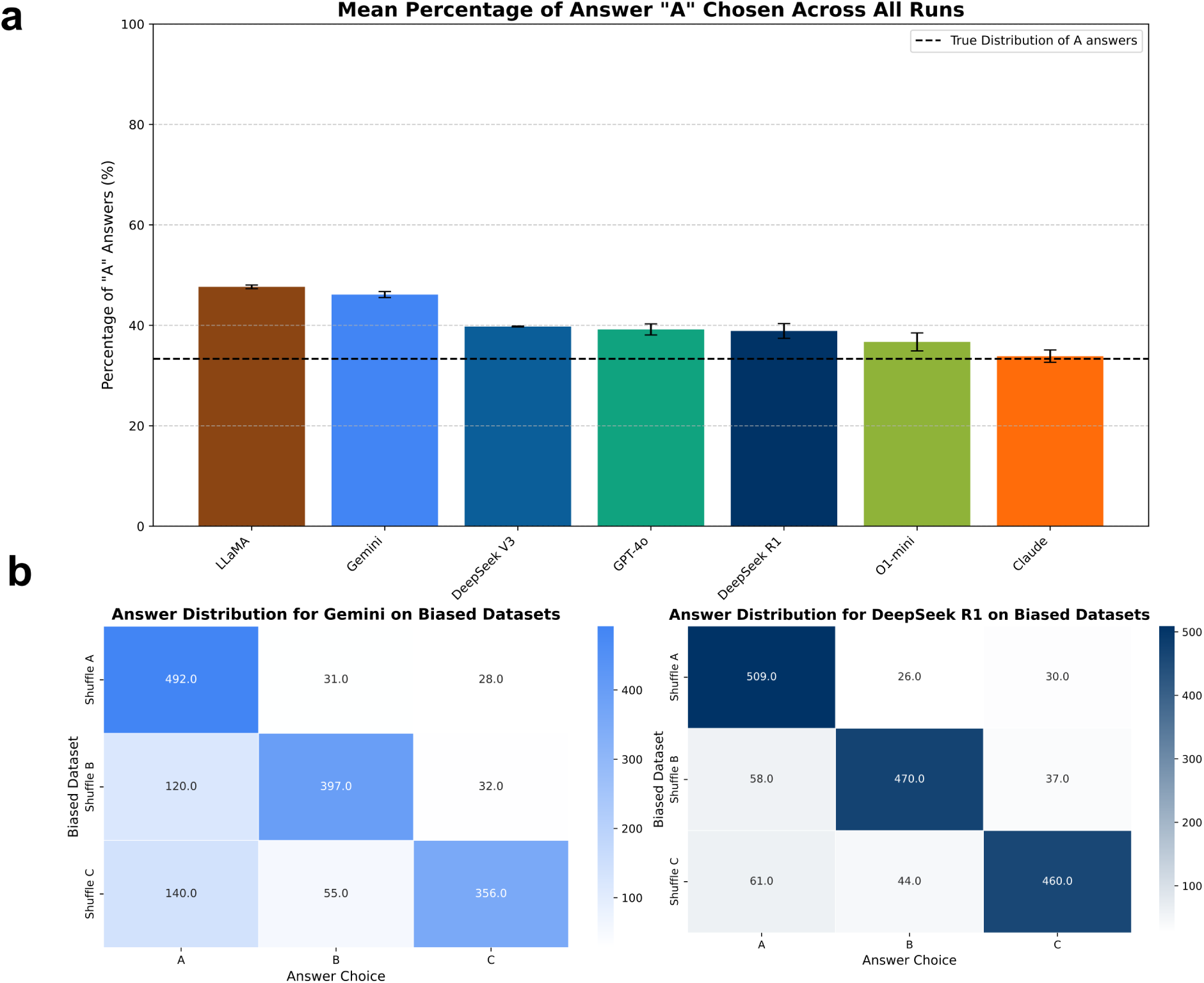
LLMs pose option bias on Expert MoBiPlant. **a)** The mean proportion of “A” responses by each LLM on the Expert MoBiPlant set over the three random shuffles. Standard deviations are denoted as error bars for each model. The distribution of correct “A” answers (%33,33) is highlighted with a dotted line. **b)** We created three alternate versions of MoBiPlant, where for each we moved all the correct answers to a specific position (the first had all correct responses transferred to “A”, the second to “B” and the third to “C”). We show the distribution of answers for these biased datasets for Gemini (left) and DeepSeek R1 (right).

### 2.5 Expert Reviews Uncover Key Gaps In LLM-generated Answers

To perform a more in-depth analysis of LLMs strengths and failure modes in plant molecular biology understanding, we prompted the models to produce open-ended answers to the Expert MoBiPlant set. Performance was assessed based on a set of criteria inspired by previous LLMs human evaluation methodologies in the medical domain^26,27^ (see Methods 5.5 for a detailed description of the human annotation procedure).

The open-ended design provides critical insights that structured benchmarks might obscure: it reveals how models construct biological arguments rather than simply selecting answers, exposing tendencies toward hallucination in underspecified scenarios and latent biases in phylum knowledge.

After the Consortium had reviewed the outputs, we created a numeric mapping that assigned each possible review a value from 0 to 100, where the values closer to zero indicate bad habits (e.g. hallucinated content) and the values near 100 indicate good outcomes (e.g. no hallucinated content in answer) on a given criterion (Figure 6.a). Remarkably, all models exhibit strong reading comprehension and maintain high topical relevance. Evidence of reading comprehension ranges from 74.0% (Gemini) to 80.6% (Claude), demonstrating that these models reliably interpret specialized prompts and extract critical details. Likewise, presence of irrelevant content scores falls between 69.1% (DeepSeek R1) and 78.2% (Claude), indicating that most generated text remains on point (higher scores indicate less irrelevant content). Potential species bias is similarly low, with values from 80.1% (DeepSeek R1) up to 83.6% (Claude), signifying broadly applicable knowledge across taxa on MOBIPLANT (higher scores indicate less species-bias).

**Figure 6.**
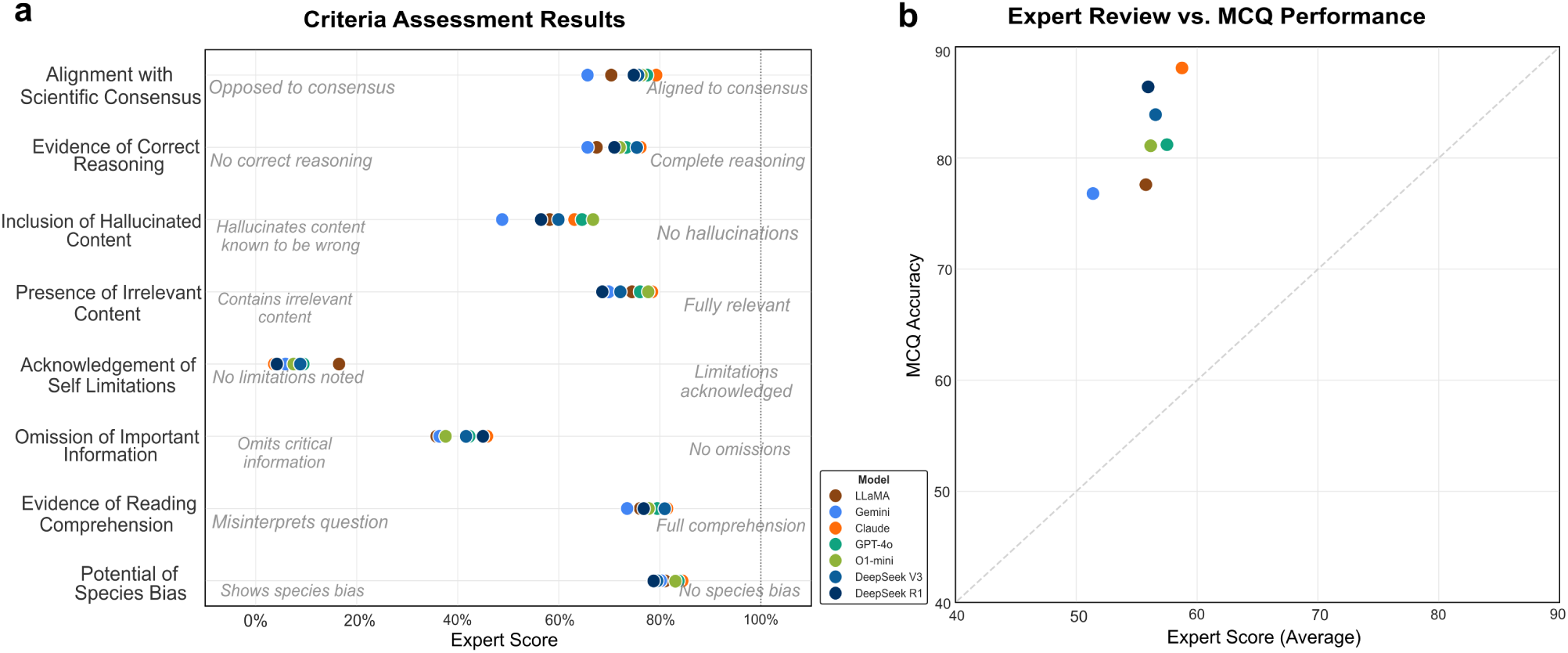
Expert evaluation on open ended answers. **a)** Human-judged scores across eight evaluation criteria. Each dot represents a model’s score (0–100%) on one criterion, with higher values indicating more positive performance (e.g., more complete reasoning, fewer hallucinations). **b)** Model’s mean human-evaluation score (x-axis, macro averaged across criteria) against its automatic evaluation accuracy (y-axis). The Pearson correlation for these scores is 0.71, demonstrating positive correlation between automatic and human-based assessments (p-value: 7.5 × 10^−2^).

In contrast, performance on factual alignment, reasoning, hallucination avoidance, and information completeness remains more moderate. Alignment with scientific consensus spans 65.7% (Gemini) to 79.3% (Claude), showing occasional deviations from established plant-science consensus, where applicable. Evidence of correct reasoning ranges from 65.7% (Gemini) to 73.9% (Claude and DeepSeek V3), indicating that causal or logical justification remains imperfect. Models score between 48.8% (Gemini) and 66.8% (o1-mini) on inclusion of hallucinated content—i.e., the proportion of responses free from fabricated claims—and only 35.8% (LLaMA) to 45.8% (Claude) on omission of important information, meaning critical details are frequently missing (lower scores indicate more omission of important information).

Remarkably, self-awareness is uniformly low across alternative LLMs: acknowledgement of limitations ranges from just 4.2% (Claude) to 16.5% (LLaMA). Interestingly, all models scored on an interval of 4.2% to 8.3% with exception of LLaMA with a remarkable 16.5%, showing stronger self-awareness in comparison to its competitors. In scientific settings, this overconfidence risks the unqualified presentation of uncertain or speculative material, skewing and potentially harming research if used without caution.

Although based on different scoring frameworks, we compared automatic evaluation accuracy based on MCQs, with human assessment scores of open-ended answers (macro-averaged across criteria; Figure 6.b). The two approaches show a positive Pearson correlation of 0.71 (p-value = 0.075), highlighting both their alignment and complementary nature in evaluating model performance. Moreover, complementary evaluations gain prominence when judging LLM’s capabilities within the plant biology domain. We show that additional axes of assessment, when conducted by experts within the field, discover significant defects of chat assistants that often go unnoticed when employing automated-only evaluation rubrics, such as MCQ.

### 2.6 Persistent Misconceptions and Hallucinations Limit LLMs In Plant Science Applications

Several notable failure modes emerged during open-ended assessments of model performance in plant biology (Table 1). *Cross-domain bias* was observed, with models incorrectly applying mechanisms from animal or human biology to plant systems. *Species confusion* occurred when models conflated traits or gene functions across distinct plant species, leading to biologically inaccurate generalizations. This is common when extrapolating well characterized mechanisms in model organisms (e.g. Arabidopsis thaliana) to other species. Errors in *acronym interpretation* were also common, where plant-specific genes were mistaken by completely different genes but similar in characters. Additionally, *terminology gaps* were evident, misunderstanding specialized plant physiology terms. The models often relied on *outdated knowledge*, failing to incorporate recent advances in emerging areas. In several instances, models *hallucinated molecular pathways*, combining unrelated pathways or involving incorrect genes in a given molecular mechanism. Finally, there was a marked *environmental misunderstanding*, with the models struggling to reason about field conditions, stress responses, such as the complex temperature- or drought-related physiological adaptations. Collectively, these failure patterns highlight key challenges for the accurate deployment of LLMs in plant-specific biological contexts.

**Table 1.**
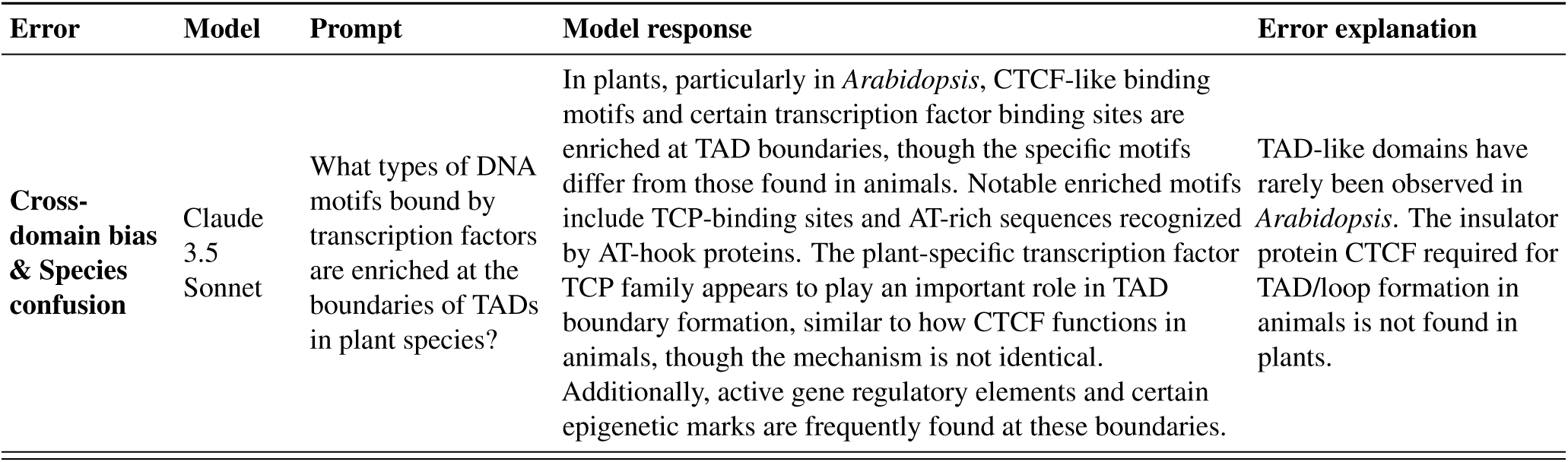
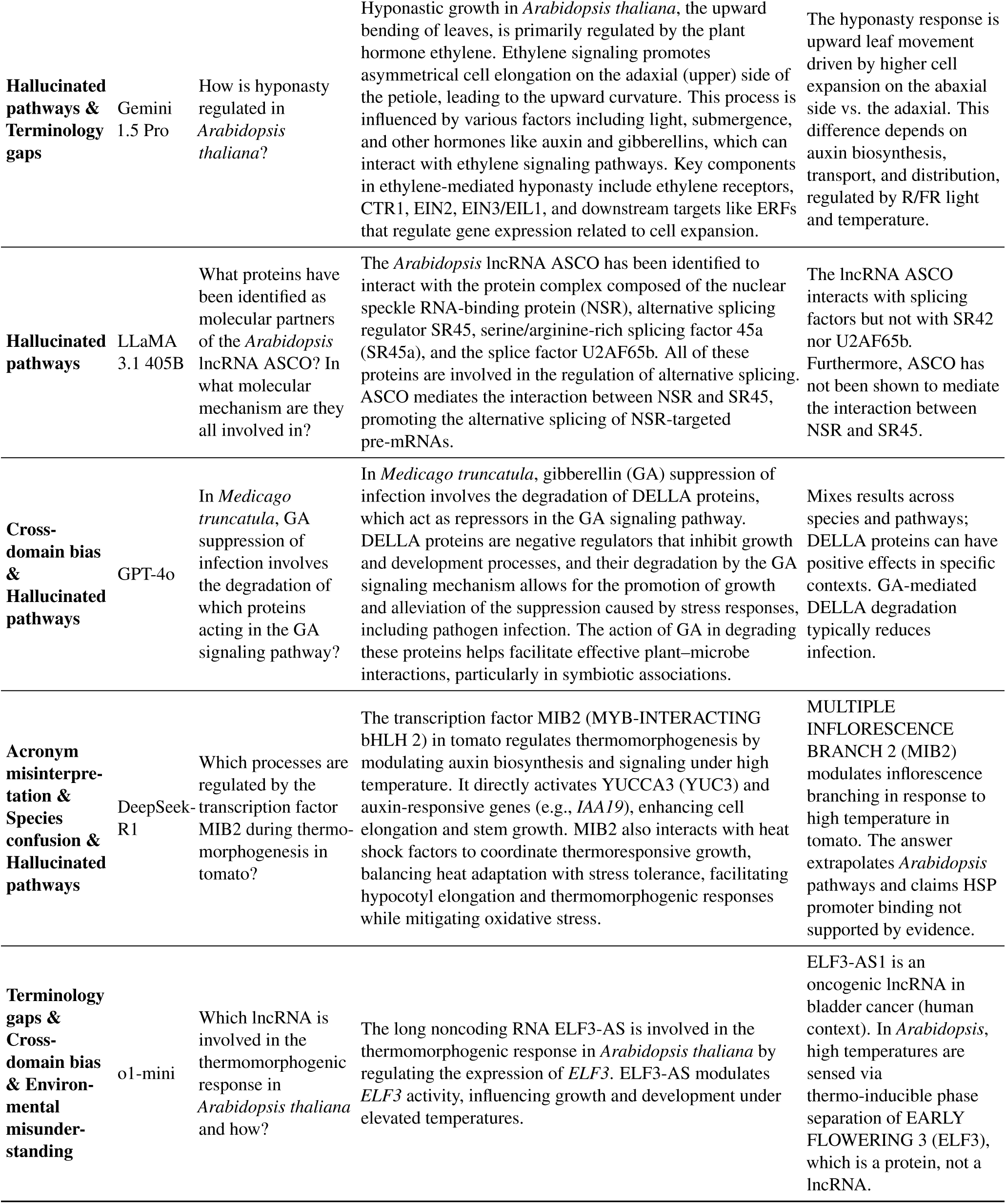
Examples of common failure modes sighted on the open-ended assessment of Expert MoBiPlant. Additional examples can be examined in Supplementary Table 3.

It has been previously reported that LLMs often struggle to provide accurate references when responding to scientific queries^28^, such as correctly identifying citation attributes like Title or Authors. In the Expert MoBiPlant set, members of the Consortium included the correct reference corresponding to the most reliable article addressing each question. During the evaluation phase, we prompted each LLM to provide a supporting reference for its answer. Although these references were reviewed by experts to assess their accuracy, the diversity of errors made a systematic evaluation challenging. In most cases, the DOI and title provided by the models were either incorrect or entirely fictitious. As illustrated in Supplementary Table 4, even for the most cited article in the dataset—and accordingly, showing remarkably good performance by LLMs delivering the related reference—the articles referred by the models exhibited various types of errors. These included: (i) the correct title paired with an unrelated DOI; (ii) the correct title with a non-existent DOI; (iii) a fabricated title with a fabricated DOI; among other misleading combinations.

### 2.7 LLM agents lag behind specialized DNA language models in plant DNA tasks

MoBiPlant assesses LLMs as conversational systems operating on natural language. Yet an increasingly relevant question is how these models compare to biological encoder-decoder models when the task requires understanding DNA sequences. To address this, we evaluated ChatNT^12^, a multimodal architecture that combines a natural language encoder with a DNA encoder module, producing outputs by jointly processing sequence and textual inputs (MOBIPLANT results in Supplementary Figure 2). Unlike the chat-based LLMs evaluated so far, ChatNT was directly fine-tuned on each target task. As a counterpart, we included two recently released agentic models; GPT-5.2^29^ and Claude Sonnet 4.6^30^ (MOBIPLANT results in Supplementary Figure 2). Both models were prompted as AI agents with access to web search, allowing them to fetch external information to inform their responses. To construct an evaluation set, we took the plant DNA datasets reported in the ChatNT paper and selected those involving plant species (see Methods 5.6).

Remarkably, ChatNT surpassed both agentic models on every task, despite operating at a fraction of the parameter count. On the classification benchmarks (Figure 7.a), ChatNT achieved 77.7% accuracy on lncRNA detection and 73.8% on enhancer detection, compared to 51.4% for both GPT-5.2 and Sonnet 4.6 on the lncRNA task and 56.1% for GPT-5.2 on enhancer detection. Notably, Claude Sonnet 4.6 refused to answer all enhancer detection queries due to refusal triggers. On the regression tasks (Figure 7.b), ChatNT achieved substantially lower mean squared error than either agentic model across both *Nicotiana tabacum* leaf (MSE = 0.894 vs. 14.954 and 4.430) and *Zea mays* protoplast (MSE = 0.610 vs. 10.924 and 5.603) tasks. These results suggest that a dedicated DNA encoder, along with direct fine-tuning on the target task, provides a decisive advantage for sequence-level plant biology tasks. Conversely, although GPT-5.2 and Sonnet 4.6 can leverage web retrieval to access relevant literature, this does not compensate for the absence of explicit sequence modelling capacity, pointing to a fundamental limitation of general-purpose language models when confronted with tasks that require interpreting information at the nucleotide level rather than knowledge retrieval queries.

**Figure 7.**
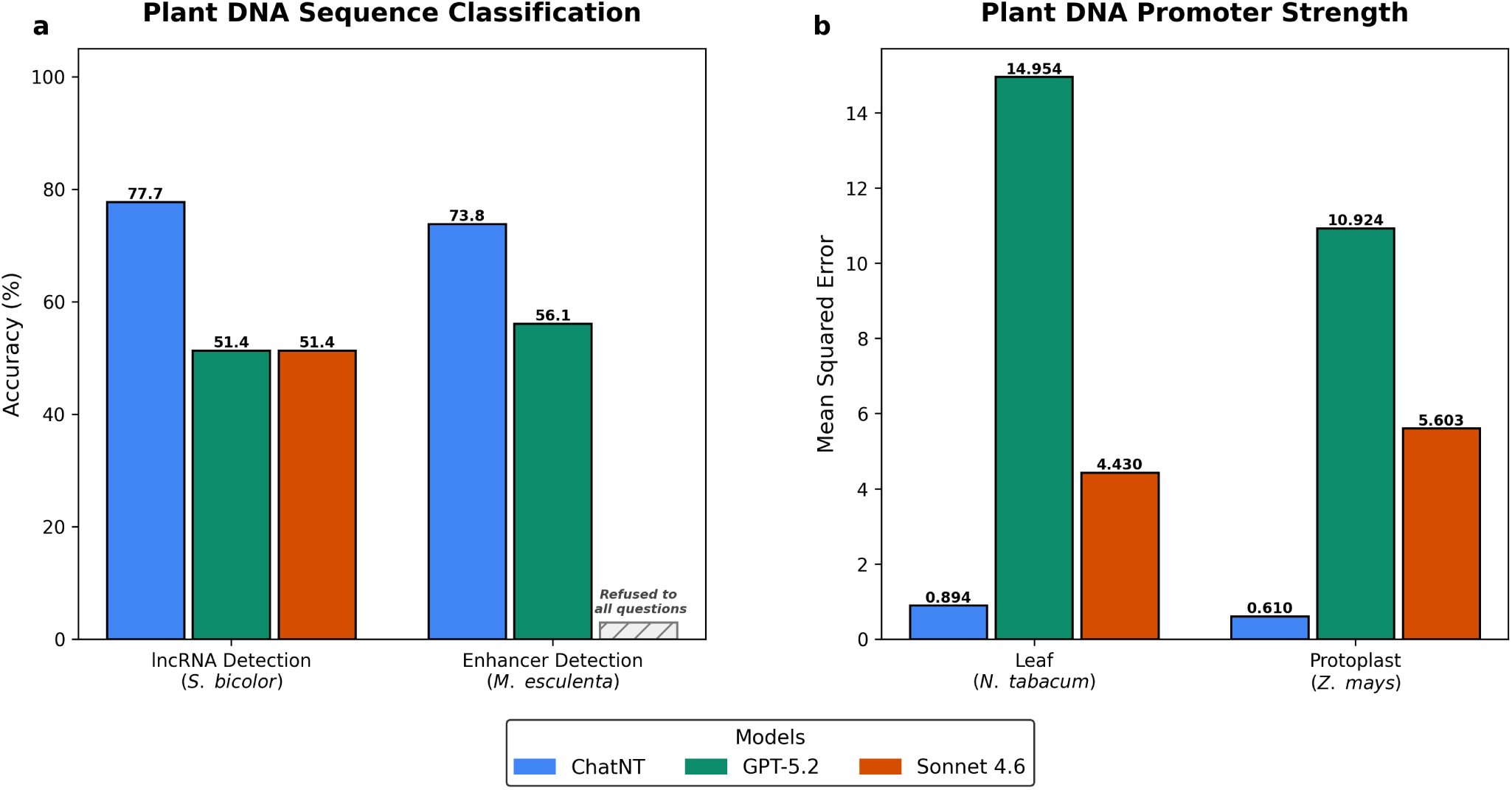
ChatNT outperforms general-purpose LLMs on plant DNA sequence tasks. **a)** Classification accuracy of ChatNT, GPT-5.2, and Claude Sonnet 4.6 on two plant DNA sequence classification benchmarks: lncRNA detection in *Sorghum bicolor* and enhancer detection in *Manihot esculenta*. Claude Sonnet 4.6 refused to answer all questions in the enhancer detection task (hatched bar). **b)** Promoter strength prediction error (mean squared error, MSE) of the same three models across two plant tissues: *N. tabacum* leaf and *Z. mays* protoplast. Lower MSE values indicate better predictive performance.

To further assess whether LLMs can interpret biological sequence information, we designed a series of structured query pipelines (Supplementary Figure 3) in which models were prompted to infer gene identity and species of origin directly from DNA sequences, including both complementary/copy DNA (cDNA) and genomic inputs. We selected representative gene sequences from *Arabidopsis thaliana*: *PIF4*, *AGAMOUS*, *BRC1* and *WOX5*, as these genes span distinct developmental and regulatory pathways and are well-characterized, providing a controlled yet biologically diverse test set. Task complexity was progressively increased by introducing sequence perturbations, such as different types of mutations, to evaluate robustness under more challenging conditions. For structured evaluation on these queries, we developed a rubric for scoring model responses and reasoning traces (see Methods 5.7). We found that across all pipelines, ChatGPT-5.2 and Claude Sonnet 4.6 failed to reliably recover even the gene identity (Supplementary Figure 3.a). Correct identification was limited to a single instance (*AGAMOUS*) by Claude (Supplementary Table 5). Interestingly, Claude also correctly suggested that the queries corresponded to genes encoding transcription factors, although the genes identified were mostly wrong. Moreover, Claude showed improved performance, occasionally inferring the correct species (*Arabidopsis*). Although model outputs frequently exhibited internally coherent, stepwise reasoning, this apparent reasoning did not translate into accurate predictions. Together, these results reveal a fundamental disconnect between linguistic reasoning and biological sequence comprehension, underscoring current limitations of LLMs in extracting meaningful information from DNA. Importantly, because ChatNT answered in the classification/regression task format in what it was instruction fine-tuned, a direct comparison with ChatNT was not feasible in this dataset.

## 3 Discussion

### 3.1 Domain-specific Benchmarking Is Critical For Assessing LLM Performance

As mentioned above, LLMs have been extensively evaluated in several high-stakes domains, with the medical field being one of the most prominent examples^26, 27^. In fact, with domain-specific training, LLMs have reached or even surpassed the passing thresholds on multiple-choice exams designed for human medical students, demonstrating their ability to retrieve and reason over well-established, textbook-based knowledge. However, the situation is markedly different in the domain of plant molecular biology. While basic concepts in botany or elementary plant molecular biology may be found in standard textbooks (somewhat analogous to consolidated anatomical knowledge in medicine), the majority of relevant and up-to-date information resides in scientific literature, such as peer-reviewed research papers. This calls for a dedicated, domain-specific evaluation frameworks in plant molecular biology like MoBiPlant. Accurately assessing LLM performance in this field requires the involvement of subject-matter experts capable of judging the model’s capacity to understand, synthesize, and apply cutting-edge research findings.

### 3.2 Automatic MCQ Evaluation And Expert Human Analysis Offer Complementary -Yet Correlated- Insights

Our results reveal that while state-of-the-art models achieve high multiple-choice accuracy (>75%) and strong comprehension in plant biology (74–81 scoring points), they still suffer from moderate factual misalignment (66–79 scoring points), frequent hallucinations (49–67 scoring points), and poor self-awareness (<17 scoring points). Human evaluations correlate positively with automatic MCQ scores while core model behaviours are addressed: open-ended judgments of reasoning, omission, and content relevance are substantially lower (Figure 6.b).

MCQ formats provide straightforward accuracy measures but can mask potential hallucination, lack of self-awareness and model biases. Additionally, uncovered failure modes highlight the risk of open-ended queries. Recent work shows that MCQ evaluation often underestimates open-ended capabilities and suffers from “first-token” misalignments between predicted option letters and actual text answers^31^. Moreover, relying solely on MCQ for evaluation risks “fake alignment” where high scores may hide poor factual grounding when models generate unconstrained text^32^. The experiment conducted on the previous section, where we show models election bias on plant molecular biology questions, sets the need for robust supporting evaluation frameworks.

Automated metrics moderately correlate with human judgments in knowledge-intensive settings. In our plant-biology domain, models are clustered by MCQ performance but scattered widely in expert criteria (hallucinations, reasoning, omissions), underscoring the need for complementary human assessments or more nuanced automated proxies. Moreover, we show that human evaluation provides richer and complementary information to that obtained through MCQ benchmarks, as models with similar MCQ scores can differ by 10–20 scoring points in expert-judged performance when it comes to assessments of reasoning quality, omission rates, and content relevance for example. A natural question is what reference performance our accuracy values should be compared against. The 565 Expert MoBiPlant questions were authored by 112 specialists each contributing items from their own subfield, so the benchmark reflects the collective knowledge of the Consortium rather than what any individual scientist (or even a PhD candidate given adequate time and literature access) could reasonably achieve. Consequently, 100% accuracy is therefore not a meaningful target for human experts, and the values reported here should be interpreted as performance relative to expert consensus. Establishing an empirical human baseline for comparison would require a dedicated study design controlling for expertise, time, and literature access, which we view as an important direction for future work.

### 3.3 LLM Understanding Scales With Paper Citations

We observe a pronounced uplift in model accuracy for questions derived from highly cited sources (501–1,702 citations) versus sparsely cited works—paralleling findings that over-represented data drive LLM competencies^33^. These effects hint at the influence of training-data composition: LLMs internalize frequent patterns but struggle with under-represented or recent material.

Crucially, performance is heavily skewed by training-data biases: by utilizing Synthetic MoBiPlant we show that questions originated from review articles consistently pose higher accuracy scores than those from primary research articles (Figure 4), highlighting the fact that review articles discuss well-established ideas within the field that are more likely to be reflected in the models’ training data and thus more readily recognized.

Although general species-bias scores remain high (80–84%, high scores indicating low bias), critical omissions (36–46%) and hallucinations (49–67%) indicate uneven coverage of specialized plant-biology knowledge. Domain benchmarks like BioLLMBench confirm similar gaps in bioinformatics tasks^34^, motivating targeted corpus expansion.

### 3.4 Hybrid Multi-agent Systems For Plant Molecular Biology

General-purpose LLMs demonstrate substantive command of plant molecular biology concepts, as evidenced by strong MCQ performance and coherent open-ended responses on MOBIPLANT, yet fail systematically when tasks require reasoning at the nucleotide level, even when equipped with agentic capabilities. Recent domain-augmented systems such as PlantScience.ai^35^ (which couples a general-purpose LLM with a graph-based plant biology knowledge base) represent a meaningful advance for factual retrieval and citation-grounded responses, but remain confined to the natural language domain and offer no capacity for biological sequence analysis. ChatNT inverts this: its dedicated DNA encoder confers an advantage on sequence-level tasks, outperforming agentic models across a wide variety of classification and regression benchmarks, but instruction-finetuning renders it less capable of use in DNA tasks different from those in the training dataset. Overall, the results indicate that it is still an open and worthwhile question whether hybrid architectures can support both natural language reasoning and nucleotide-level inference without compromising either capability.

An alternative design may consist of a multi-agent framework in which a general-purpose LLM acts as an orchestrator which can handle task decomposition, contextual reasoning, and result interpretation, delegating sequence-level inference to specialised biological foundation models such as DNABERT^36^, Evo2^37^ or ESM^38^ via structured tool calls. Such a design should preserve the conversational and reasoning capabilities of the LLM without subjecting it to fine-tuning distributions that would degrade its generality, while simultaneously exploiting the sequence modelling capacity of models trained explicitly on genomic or proteomic data.

## 4 Conclusions

LLMs are increasingly being used by a broad audience—from high school and undergraduate students to early-career and established researchers. While their applications continue to expand, the tendency to rely on LLMs as a substitute for critical document-based research—receiving synthesized answers presented as factual—poses significant risks. In the field of plant molecular biology, in particular, careful validation of LLM-generated content is still needed to prevent the dissemination of misconceptions and inaccurate information.

Our comprehensive benchmarking reveals that while current LLMs excel at parsing and responding to structured plant biology prompts—achieving high MCQ accuracy and strong reading comprehension—they consistently struggle with deeper scientific rigor and self-awareness. Factual alignment remains only moderate, hallucinated content persists at nontrivial rates, and essential details are often omitted. Crucially, all models exhibit a pronounced inability to recognize or communicate their own uncertainties, risking the unqualified presentation of speculative or incomplete information. Interestingly, while ChatGPT (particularly GPT-4o) is by far the most widely adopted model among researchers in our consortium—as reported in our survey of preferred assistants—it does not consistently rank as the top performer in our benchmark evaluations. Although it achieves solid results in both MCQ and open-ended formats, models such as Claude 3.5 Sonnet and DeepSeek R1 tend to outperform it across key evaluation dimensions. Furthermore, Claude 3.5 Sonnet and O1-mini were the only LLMs assessed that did not exhibit any bias towards option A in MCQs.

Our findings underscore the limitations of evaluating LLMs solely on closed-form tasks and highlight the value of expert human assessment in revealing hidden failure modes. To advance toward trustworthy scientific assistants, it will be crucial to integrate specialized biological foundational models alongside retrieval augmentation or search tools in LLM-based agentic workflows to ground outputs in authoritative plant databases, applying domain-specific fine-tuning to reinforce structured knowledge, and incorporating uncertainty calibration mechanisms that enable models to flag low-confidence assertions. Our comparison between ChatNT and agentic LLMs on plant DNA sequence tasks illustrates this point concretely: despite having access to web retrieval, general-purpose models were markedly outperformed by a model that integrates a dedicated DNA encoder and was fine-tuned on the target task, suggesting that for sequence-level reasoning, architectural specialization currently provides an advantage that retrieval alone cannot bridge.

By adopting this multi-pronged strategy, future models can not only bolster their factual accuracy and completeness but also develop the capacity necessary for safe and reliable deployment in research scenarios. The path forward lies in blending rigorous evaluation frameworks with targeted architectural enhancements to realize the full potential of chat assistants in plant biology.

## 5 Methods

### 5.1 Dataset Design and Curation

The construction of the MoBiPlant Benchmark involved the creation of a consortium of 112 researchers in plant molecular biology (the MoBiPlant Consortium), selected to represent diverse geographical regions, career stages, institutional affiliations, and demographic backgrounds. Contributors spanned disciplines encompassing molecular mechanisms (gene regulation, genome and genomics, cell biology and signaling), organismal processes (growth and development, hormones, physiology and metabolism, evolution), and applied domains (biotechnology, environment). Each expert authored five multiple-choice questions (MCQs) aligned with their specialization, adhering to rigorous guidelines ensuring question clarity, plausibility of distractors, and balanced option lengths to mitigate selection biases. A complete protocol detailing question design principles is provided in Supplementary Material Section H.

To elevate the dataset size, we further boosted question count using a frontier LLM which was prompted to generate 1075 additional question-answer pairs from human-selected papers. We detail the extraction, processing and creation in Section 5.2.

Each question in MoBiPlant includes rich metadata detailing the plant species involved, the specific sub-discipline it relates to, and the source publication where the correct answer can be verified (with the exception of eleven more general questions, where their authors claimed that multiple sources where required). These fields enhance the utility and transparency of the benchmark by enabling topic-based filtering, ensuring traceability to original sources, and offering a clearer view of the content’s scope. Figure 2.b-d presents the distribution of these attributes, illustrating the representation of species, topic areas, and source diversity across the dataset.

### 5.2 Synthetic Data Generation

The production of synthetic datasets has been of uprising use over the recent years^39–45^, with primarily focus on model distillation and model evaluation. In this case, we propose a framework for systematically extracting multiple-choice questions from a curated set of manually selected publications within plant molecular biology throughout the years 2001 to 2025.

For the selection of published papers, a group of specialists selected subsets of relevant work documents in the following journals: (1) research articles from New Phytologist, Plant Journal, Plant Physiology, The Plant Cell, Molecular Plant, Journal of Experimental Botany, and Nature Plants; and (2) review articles from Trends in Plant Science, and Current Opinion on Plant Biology (see Supplementary Table 6: MoBiPlant Journal distribution). This curation ended with a total of 215 documents (at least one per year and journal, if already existing) with fine-grained information within the field.

Consequently, we prompt a frontier LLM (Gemini 2.5 Pro) to act as an MCQ extractor when a work is given as pdf. The process is similar to what is solicited to the human collaborators: generate 5 multiple-choice questions with one correct answer and two incorrect answers (distractors). The instructions intend to be as similar as possible to the guidelines prompted to humans (see Supplementary Material Section I). The idea was to ensure a clear and unambiguous question that is directly linked with a unique fact or conclusion from the given work, with its respective correct answer and distractors generated by tweaking the information on the correct answer in a way that keeps semantic and terminology-related feasibility. In addition, the instructions include the metadata tagging of each question (whether there is a plant species involved in the query and its respective area), ensuring compatibility with the human-generated MCQ. By generating 5 questions for every one of the 215 studies collected on the previous step, we ended up with 1075 additional multiple-choice questions tagged and compatible with the previous format, which constitute the Synthetic MoBiPlant dataset.

For the validation of this methodology, we randomly sampled 20 questions ensuring equal representation from all journals for human revision, which confirmed the suitability of the generated questions. This was performed by the same experts that proposed the studies.

### 5.3 Model Selection

To conduct this evaluation, we selected a mixture of open and closed models, excelling in several general benchmarks^23, 46, 47^ and validated by the MoBiPlant Consortium top chat-assistant choices (Figure 1.g). There are several biologically tuned language models centered in biomedical knowledge^48–51^, but they are comparatively small to the models widely adopted as first-hand large chat assistants by the community. This is why we focused on benchmarking the following selection comprising GPT-4o^17^, Claude 3.5 Sonnet^22^, Llama 3.1 405B^21^, Gemini 1.5 Pro^16^ and Deepseek V3^19^ as well as recently released reasoning models, featuring o1-mini^18^ and Deepseek R1^20^.

All models were instantiated with a value of 0.7 for temperature and a max generation length of 4096 tokens to ensure an equal evaluation environment and mimic that of assistant usage settings.

### 5.4 Automated MCQ Evaluation

To extensively assess model performance, we employ two parallel evaluation designs: Automated MCQ evaluation and Human Evaluation (see next section for a detailed description of the Human Evaluation Protocol).

For automatic evaluation we directly computed the accuracy of model answers over the set of MCQ questions. This method presents models with questions and predefined answer choices, followed by automated extraction of their selections for comparison against ground-truth answers. While this approach offers advantages—such as providing a structured framework for systematic evaluation and simplifying answer extraction by constraining responses to predefined choices—it introduces three main limitations: 1) exploitation of shortcuts/biases^52^: models may exploit statistical patterns or suboptimal distractors (e.g. option order) to guess correct answers without genuine comprehension, artificially inflating performance metrics; 2) limited real-world applicability^53, 54^: MCQ formats may inadequately reflect real-world applications, where open-ended queries are the norm rather than constrained selections; and 3) choice-order sensitivity^55^: performance can vary based on the order of answer choices, introducing inconsistencies unrelated to model capability.

To address concerns of model over-reliance on statistical patterns and choice-order sensitivity, we generated three permutations of the MCQ dataset, each containing questions with randomly shuffled answer options. By evaluating model performance across all permutations, we quantified variability in accuracy attributable to option order using the standard deviation of scores between shuffled versions. This approach directly measures robustness to positional bias while ensuring reproducibility across stochastic configurations.

We designed a chain-of-thought prompt^56^ to elicit reasoning in model output generation. The prompt instructs the model to reason through the correct answer in order to enhance answer generation. This approach is natural and aligns the real-world application of MCQs, encouraging the model to generate a step-by-step reasoning before pointing to the final answer between `<answer></answer>` tags, which is later extracted using string-matching techniques. Since this methodology introduces possible parsing errors in model generation we report the valid accuracy when scoring with this method. This is, the accuracy over all examples in which a model answer could be parsed.

### 5.5 Human Evaluation

To address the limitations of automated multiple-choice assessments, we implemented a human expert evaluation protocol adapted from existing clinical knowledge language model validation frameworks^26, 27^. This approach begins with a priori definition of domain-specific evaluation criteria, developed collaboratively with plant molecular biologists to reflect field-specific priorities. Two core criteria were established in addition to the existing: (1) Species bias, assessing whether model answers overgeneralize findings beyond the queried species (e.g., attributing Arabidopsis thaliana mechanisms to non-model plants), and (2) Consensus alignment with established plant science dogmas and theories, evaluating agreement with peer-reviewed evidence while accounting for contested or evolving concepts. This resulted in a total of 8 criteria for human evaluation, as defined in Table 2.

**Table 2.** Criteria set used for human evaluation on MOBIPLANT.

| **Criterion** | **Description** | **Rating Options & Explanations** | **Score** |
| --- | --- | --- | --- |
| **Evidence of Correct Reasoning** | Assesses whether the response follows logical reasoning to arrive at a correct conclusion. | **Yes:** The reasoning is logical, well-structured, and leads to a correct conclusion.<br>**No:** The reasoning is flawed, contains logical fallacies, or does not support the conclusion. | Yes: **1**<br>No: **0** |
| **Evidence of Reading Comprehension** | Evaluates whether the response fully understands and addresses the question, including its context and nuances. | **Yes:** Fully comprehends and correctly addresses the entire question.<br>**Partially:** Shows partial understanding but misses important details.<br>**No:** Misinterprets or ignores key aspects of the question. | Yes: **1**<br>Partially: **0.5**<br>No: **0** |
| **Alignment with Scientific Consensus** | Determines if the response aligns with the widely accepted scientific understanding. | **Aligned to consensus:** Reflects the prevailing scientific view, dogmas and/or theories.<br>**Opposed to consensus:** Contradicts widely accepted scientific knowledge.<br>**No consensus:** The topic lacks a clear, widely agreed-upon stance. | Aligned: **1**<br>Opposed: **0**<br>No consensus: (don't compute) |
| **Omission of Important Information** | Checks if critical biological details are missing, which could affect the accuracy of the response. | **Yes, great biological significance:** Major omission that alters understanding.<br>**Yes, little biological significance:** Minor omission with limited impact.<br>**No:** Includes all necessary information. | Yes, great: **0**<br>Yes, little: **0.5**<br>No: **1** |
| **Inclusion of Hallucinated Content** | Identifies whether the response contains incorrect or unsupported information. | **Yes, known wrong:** Claims fabricated/incorrect information.<br>**Yes, unverifiable:** Includes unverifiable claims.<br>**No:** Based on verifiable information. | Yes, known wrong: **0**<br>Yes, unverifiable: **0.5**<br>No: **1** |
| **Inclusion of Irrelevant Content** | Determines if the response includes off-topic or unnecessary details. | **Yes:** Contains extraneous information.<br>**No:** Stays focused on the question. | Yes: **0**<br>No: **1** |
| **Potential of Species Bias** | Assesses whether the response shows bias toward or against specific species in a biological context. | **Yes:** Exhibits species bias (e.g., general claim untrue across species, or answers about a species using other species).<br>**No:** Does not contain biased claims.<br>**Not applicable:** The question does not involve species comparisons. | Yes: **1**<br>No: **0**<br>Not applicable: (don't compute) |
| **Acknowledgement of Limitations** | Evaluates whether the response recognizes uncertainties or limitations where relevant. | **Yes:** Explicitly acknowledges relevant self-limitations.<br>**No:** Omits relevant uncertainties despite evident limitations.<br>**Not applicable:** The answer does not require discussing limitations. | Yes: **1**<br>No: **0**<br>Not applicable: (don't compute) |

For species bias, evaluators flagged answers containing unsupported taxonomic extrapolations, a critical concern when employing these systems in understudied species. Alignment with scientific consensus was scored on a ternary scale: alignment (consistent with established literature), opposition (contradicting peer-reviewed evidence), or lack-of-alignment (addressing unresolved or actively debated topics). This granular scoring acknowledges the dynamic nature of plant biology, where emerging methodologies (e.g., the advention of CRISPR-based gene editing) routinely challenge paradigms. The detailed description of every criterion with the available options that were provided to the expert reviewers is included in Table 2.

For evaluating model performance in open-ended contexts mirroring real-world scientific inquiry, we generated responses to all MoBiPlant questions using the same prompts across models. Instructions explicitly directed models to provide concise, self-contained answers without supplying the predefined multiple-choice options. The idea was to simulate situations where users do not know the answers in advance, helping to reveal subtle behaviors like guessing, overgeneralizing, or missing details that MCQs often hide.

To ensure evaluator rigor, we implemented a blind workflow: model answers were anonymized, stripped of metadata (e.g., model names, question area), and randomly shuffled across questions. Crucially, since experts in one area may not have the detailed knowledge needed to evaluate work in another, each contributor evaluated responses only to their own original questions. We used the SuperAnnotate Platform^57^ to coordinate the annotation process involving the 112 researchers from the MoBiPlant Consortium.

To standardize the manual evaluation protocol, we established a normalized scoring framework. Qualitative criteria (e.g., species bias, consensus alignment) were mapped to normalized numerical scores ranging from 0 (indicating undesirable performance) to 1 (indicating optimal performance). This transformation preserves the interpretability of expert judgments while facilitating quantitative correlation analyses. For instance, responses contradicting scientific consensus were assigned a score of 0, whereas alignment with peer-reviewed evidence scored 1; ambiguous or context-dependent answers received intermediate values. Detailed mappings are provided alongside the criteria description in Table 2.

### 5.6 Plant DNA Tasks Dataset Construction

We used the data published by ChatNT’s authors to generate a subset of plant molecular biology questions involving a DNA sequence. We took the plant DNA datasets reported in the ChatNT paper and selected those involving plant species. For each dataset, we drew a random 5% subsample, stratifying by class label for classification tasks and applying label-aware sampling for regression tasks to ensure a uniform target distribution in the final subset. This resulted in 37 sequences for lncRNA detection in *Sorghum bicolor*, 41 for enhancer detection in *Manihot esculenta*, 358 for promoter strength prediction in *Nicotiana tabacum* leaf, and 380 for promoter strength prediction in *Zea mays* protoplast. Sequences and task descriptions were formatted into natural language prompts and submitted to GPT-5.2 and Claude Sonnet 4.6 with web access enabled. ChatNT predictions were obtained by the data made available by the authors in their supplementary information.

### 5.7 Plant DNA Queries Dataset and Evaluation Rubric Construction

Each of the eight queries in the DNA Queries Pipeline (Supplementary Figure 3.a) was paired with a domain-expert rubric that defined what a model response had to assert in order to be considered correct. Rubrics were authored manually for each of the four target genes (*PIF4*, *AGAMOUS*, *BRC1* and *WOX5*) and stored in a single spreadsheet, with one row per (gene, pipeline-step) combination encoding the question text, the criterion description, a numerical weight (criterion_score), and where applicable the mutation class introduced into the sequence (Supplementary Table 7). For the gene-and-organism identification queries (Steps 0 and 1), the rubric was decomposed into two independently weighted sub-criteria: one verifying that the model named the correct gene, and one verifying that it identified *Arabidopsis thaliana* as the source organism, each contributing 5 points. For the remaining steps (2, 3, 4a, 4b, 5a, 5b), a single criterion worth 10 points was scored. This per-step weighting ensures that simple identification tasks and more complex biological-reasoning tasks contribute comparably (10 points each per gene per step) to the aggregate score reported in Supplementary Figure 3.b.

Two types of criteria were used. Single-statement criteria (e.g., “It states that the gene is *PIF4*”, or, for silent-mutation steps 4b and 5b, “It states that the phenotype is normal given that the mutation is silent, thus there is no impact on the protein translated”) admit only one acceptable formulation. Listed-statement criteria enumerate multiple distinct, biologically valid claims that an expert would accept as correct; these accepted statements correspond to those documented in Araport. For example, the criterion for the loss-of-function phenotype of *PIF4* includes five separately sufficient statements ranging from impaired thermomorphogenic hypocotyl elongation to delayed flowering at high temperature. For listed-statement criteria, the criterion is considered met if the model’s answer clearly satisfies at least one of the accepted statements as defined in Araport.

Scoring of each model response against its rubric was performed automatically using GPT-5.2 as a judge. To minimise the well-documented biases of free-form LLM judges, we constrained the judge to a single binary decision per criterion. Specifically, the judge was prompted with a system message instructing it to act as a strict binary evaluator and to return only a compact JSON object containing exactly one key, criterion_met, with values restricted to 0 or 1. The user message then provided (i) the original question presented to the evaluated model, (ii) the criterion (or, for listed-statement criteria, the enumerated set of acceptable statements), (iii) the candidate model answer, and (iv) the satisfaction rule appropriate to the criterion type. The judge was instructed to set criterion_met = 1 only if the model answer clearly satisfied the criterion; missing, incorrect, or ambiguous answers were scored 0.

For each evaluation, the binary judgement returned by GPT-5.2 was multiplied by the corresponding criterion_score weight, summed across the (up to two) sub-criteria of a step, and then averaged across the four target genes to produce the per-step performance values reported in Supplementary Figure 3.b.

## 6 Data and Code Availability

All datasets used in this study, Expert MoBiPlant and Synthetic MobiPlant are available via HuggingFace (manufernandezbur/MoBiPlant).The code used to process data, generate datasets and perform the analyses described in this paper is available at GitHub, including model

## 7 Ethical Considerations

This study evaluates the capabilities and limitations of large language models (LLMs) in the context of plant molecular biology using a benchmark developed by a global consortium of domain experts. The research was conducted in alignment with ethical principles of transparency, fairness, and inclusivity.

Expert Contributions and Consent: The MoBiPlant Benchmark was created with the voluntary participation of 112 researchers across 19 countries. All contributors agreed to participate and were fully aware of the project’s goals.

Diversity and Representation: The study prioritized inclusive participation, ensuring representation across gender identities, geographic regions, institutional affiliations, and career stages. Demographic information was collected and reported in aggregate form (Figure 1) to inform on dataset composition and potential biases.

Synthetic Data Use: The study includes synthetic data generated using a frontier LLM. These data were derived from publicly available scientific literature and used strictly for evaluation purposes. A human-in-the-loop validation process was implemented to mitigate the risk of embedding flawed or biased information. No copyrighted or proprietary content was used beyond what is permissible under fair use for scientific research.

Model Evaluation and Bias Awareness: Evaluated models are commercial and open-source LLMs that operate as general-purpose assistants. We acknowledge that these models may reflect biases inherent in their training data, including overrepresentation of certain species, geographic regions, or scientific paradigms. As such, our evaluation explicitly includes metrics for species bias and alignment with scientific consensus. We emphasize that LLM outputs should not be regarded as authoritative or error-free, particularly in understudied taxa or emergent research areas.

Use of AI in Scientific Assessment: We recognize the ethical implications of using AI-generated outputs in scientific settings. While LLMs offer utility in accelerating literature review and hypothesis generation, our findings underscore the continued need for expert oversight. To prevent misuse, we do not advocate for the replacement of domain expertise with AI, but rather for informed, cautious integration of these tools under appropriate validation frameworks.

Transparency and Reproducibility: All prompts, evaluation protocols, and scoring rubrics are provided in the Methods section and Supplementary Materials to support transparency and reproducibility. The MoBiPlant dataset is made freely available for academic research under an open license, enabling community-led extensions and audits.

## Acknowledgements

The FA lab is funded by AXA Research Fund, ICGEB, Agencia I+D+i, and the IRP NOCOSYM (CNRS). The EF lab is supported by Agencia I+D+i, a Human Frontiers grant, the Google Award for Inclusion Research and a Googler Initiated Grant. MFB was supported by the AI Safety Argentina Scholarship (AISAR) program. APOLO Biotech covered the computing expenses. We thank SuperAnnotate for giving us access to their annotation interface.

## Supplementary Material

## A The MoBiPlant Consortium

**Supplementary Table 1.**
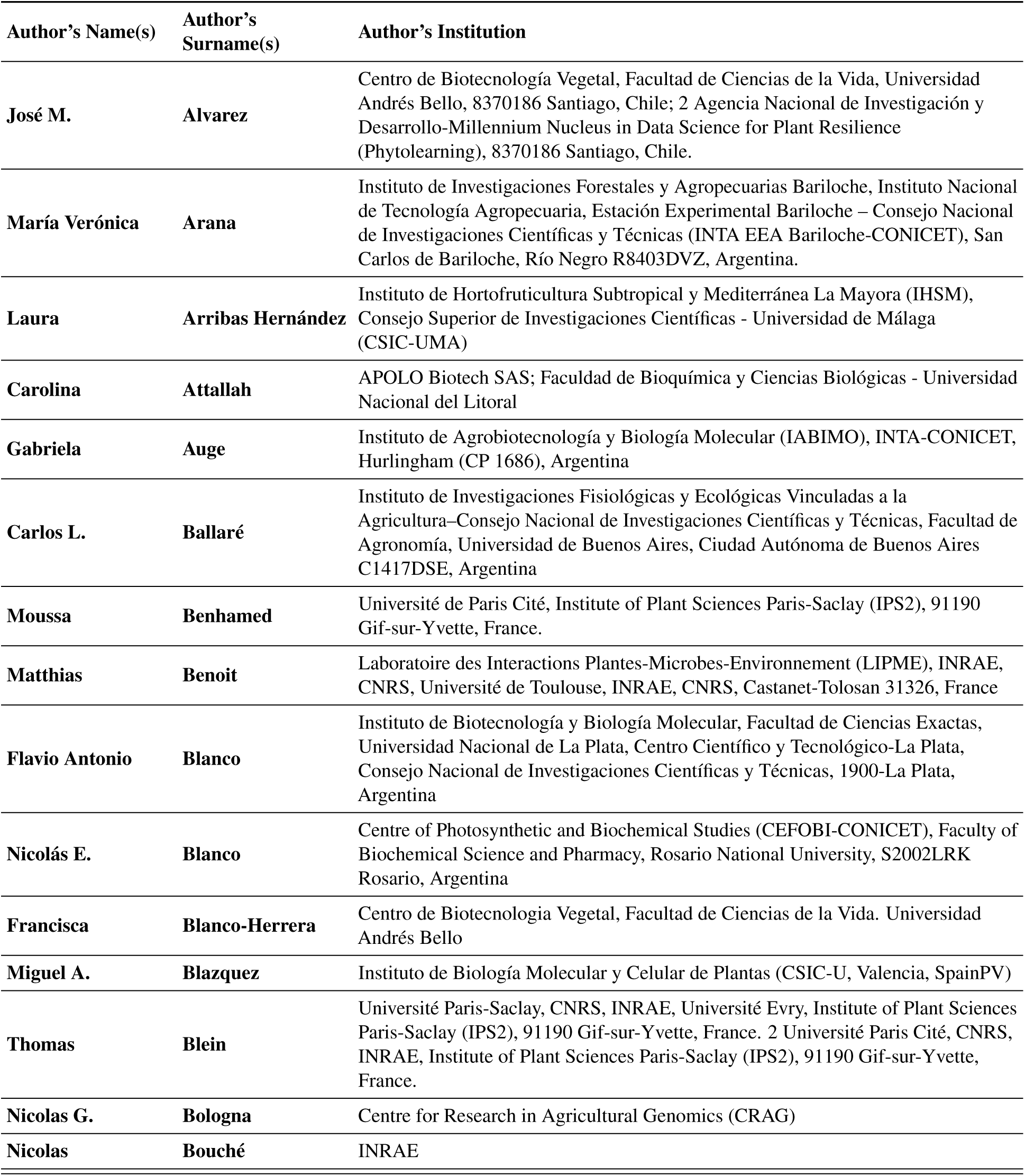
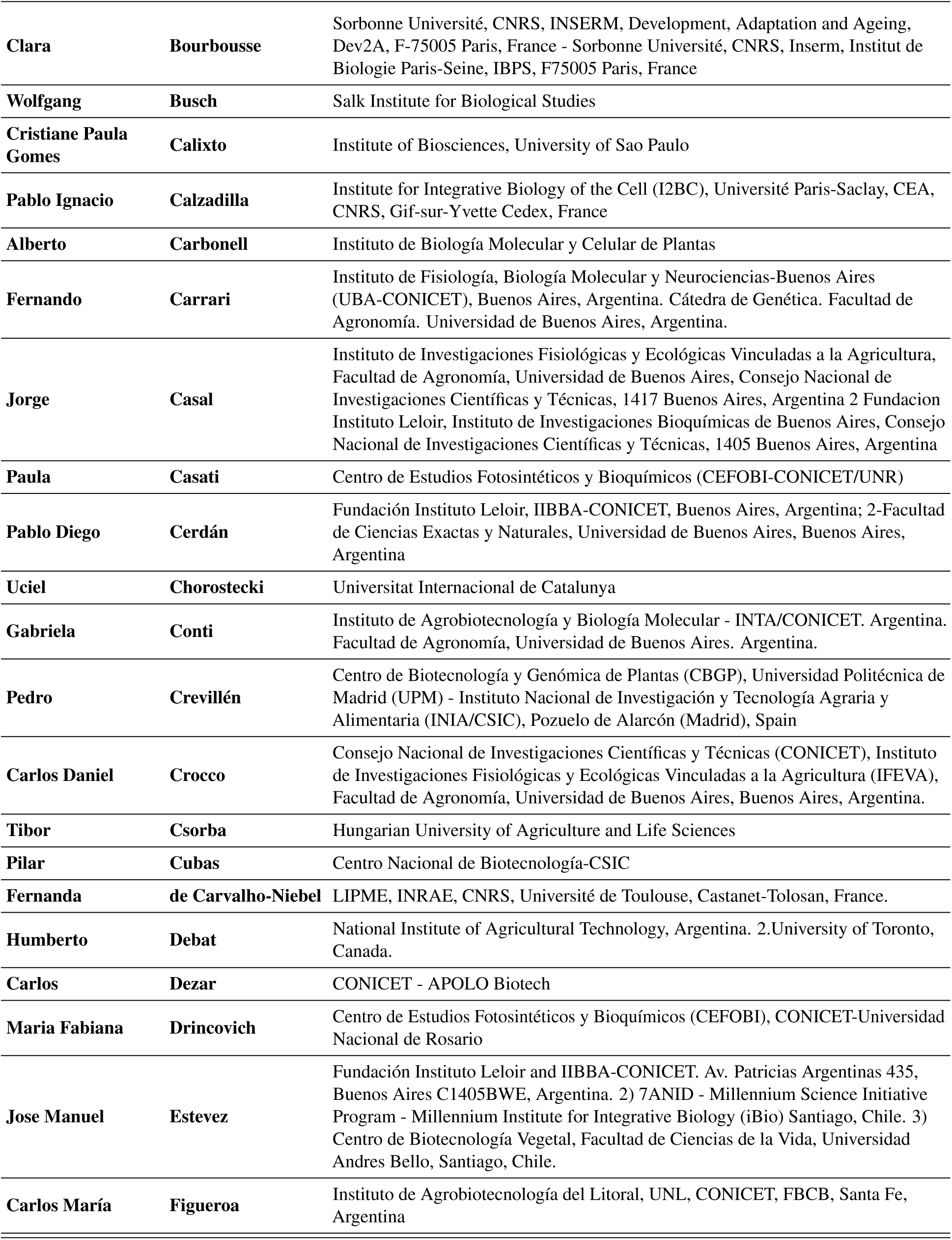
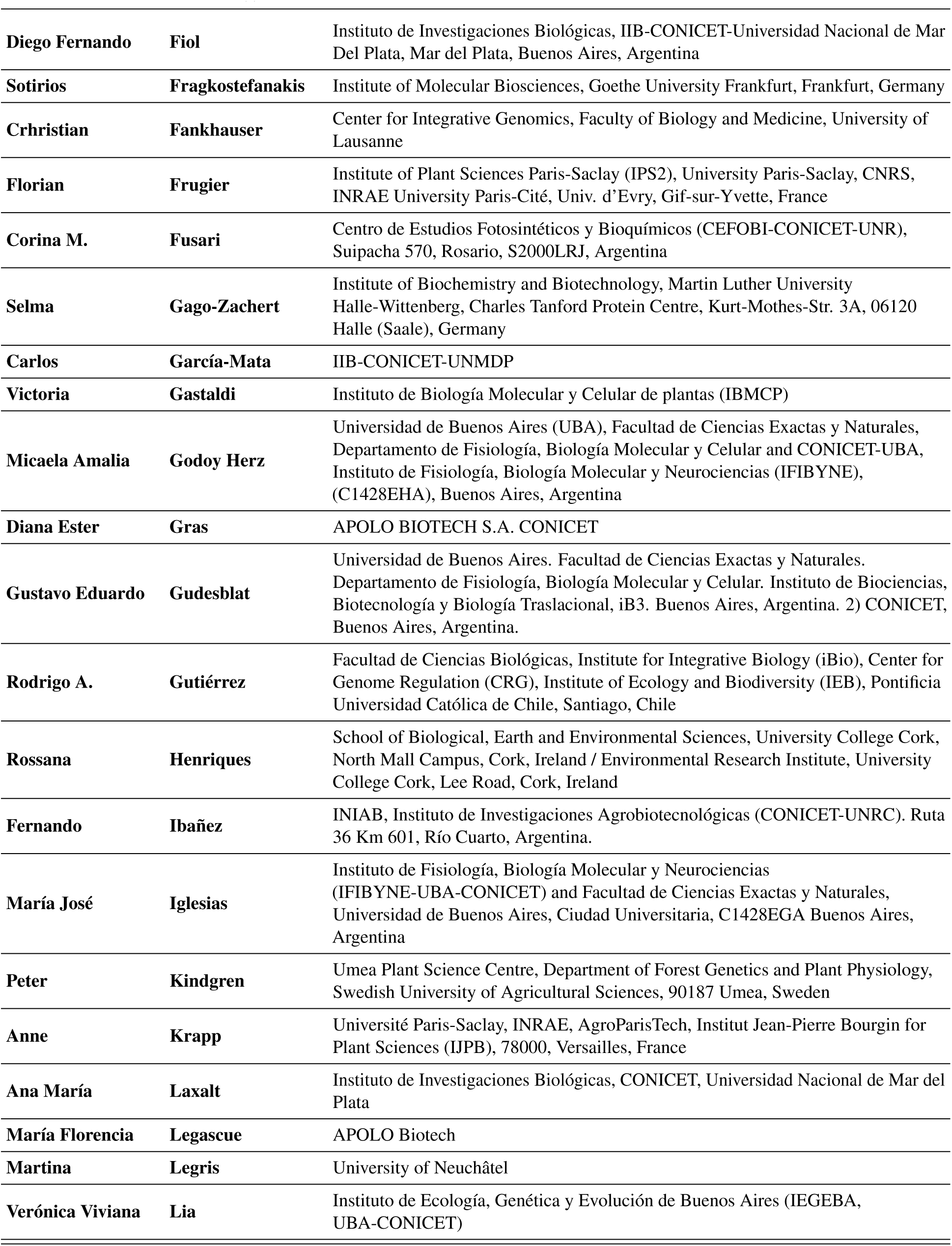
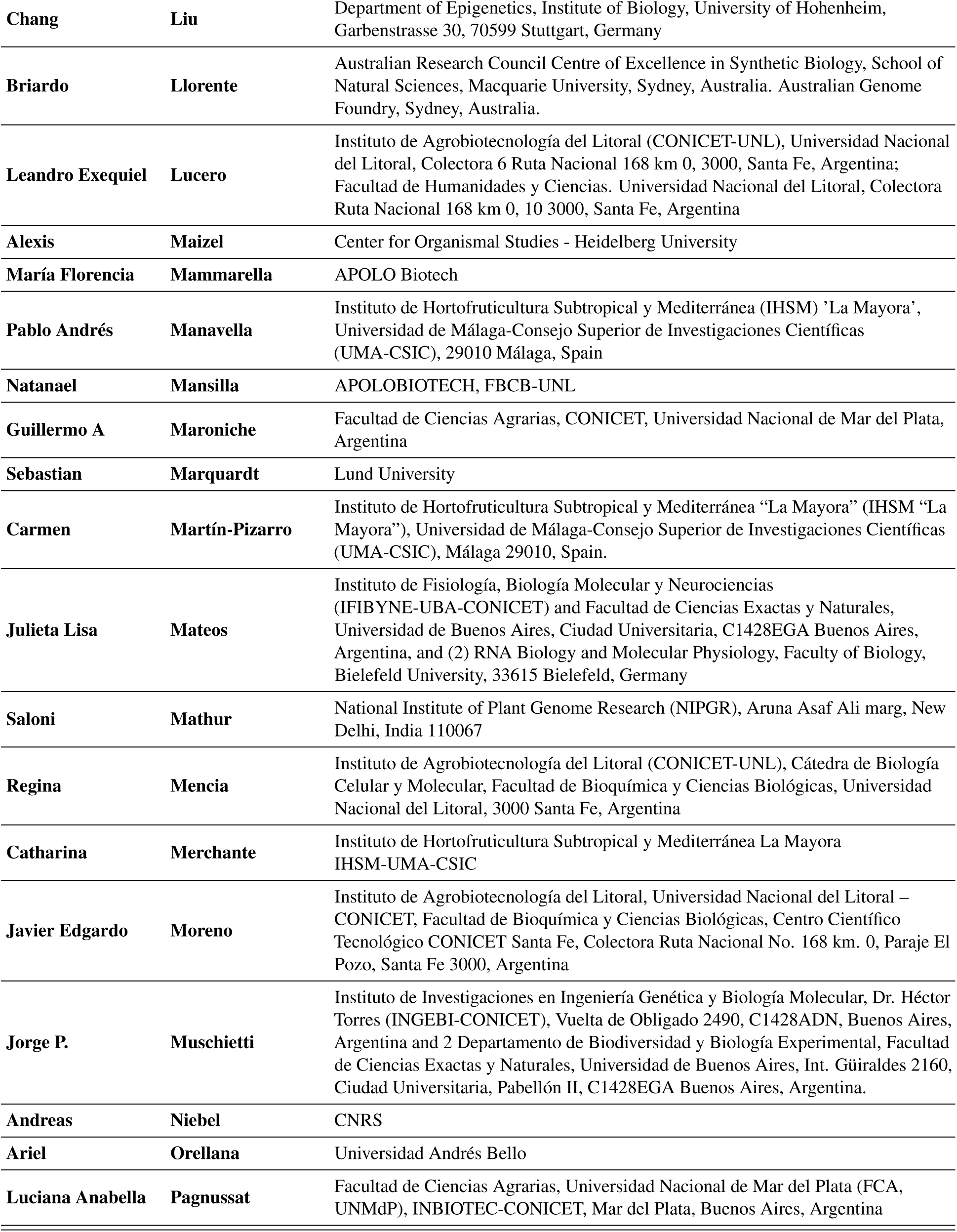
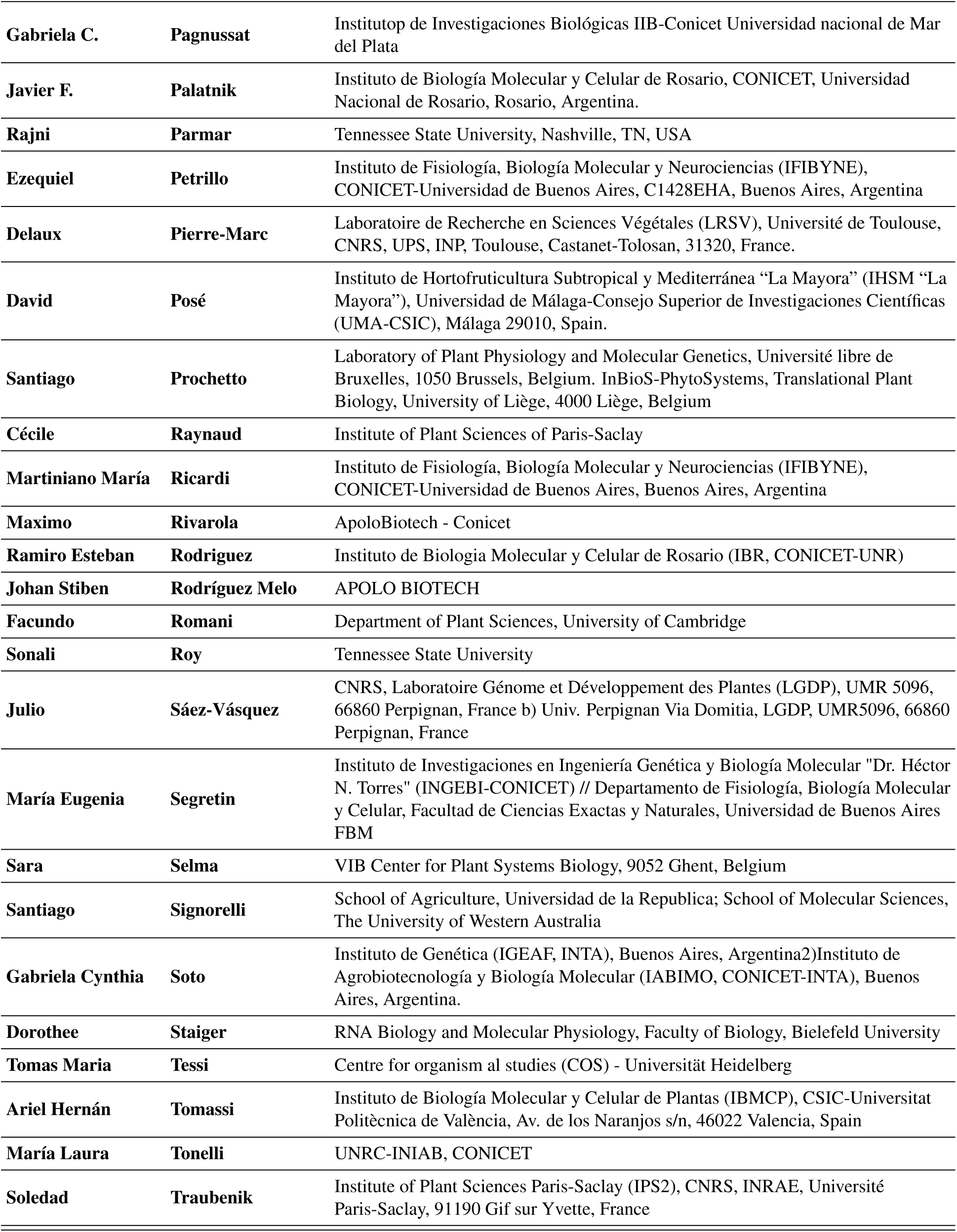
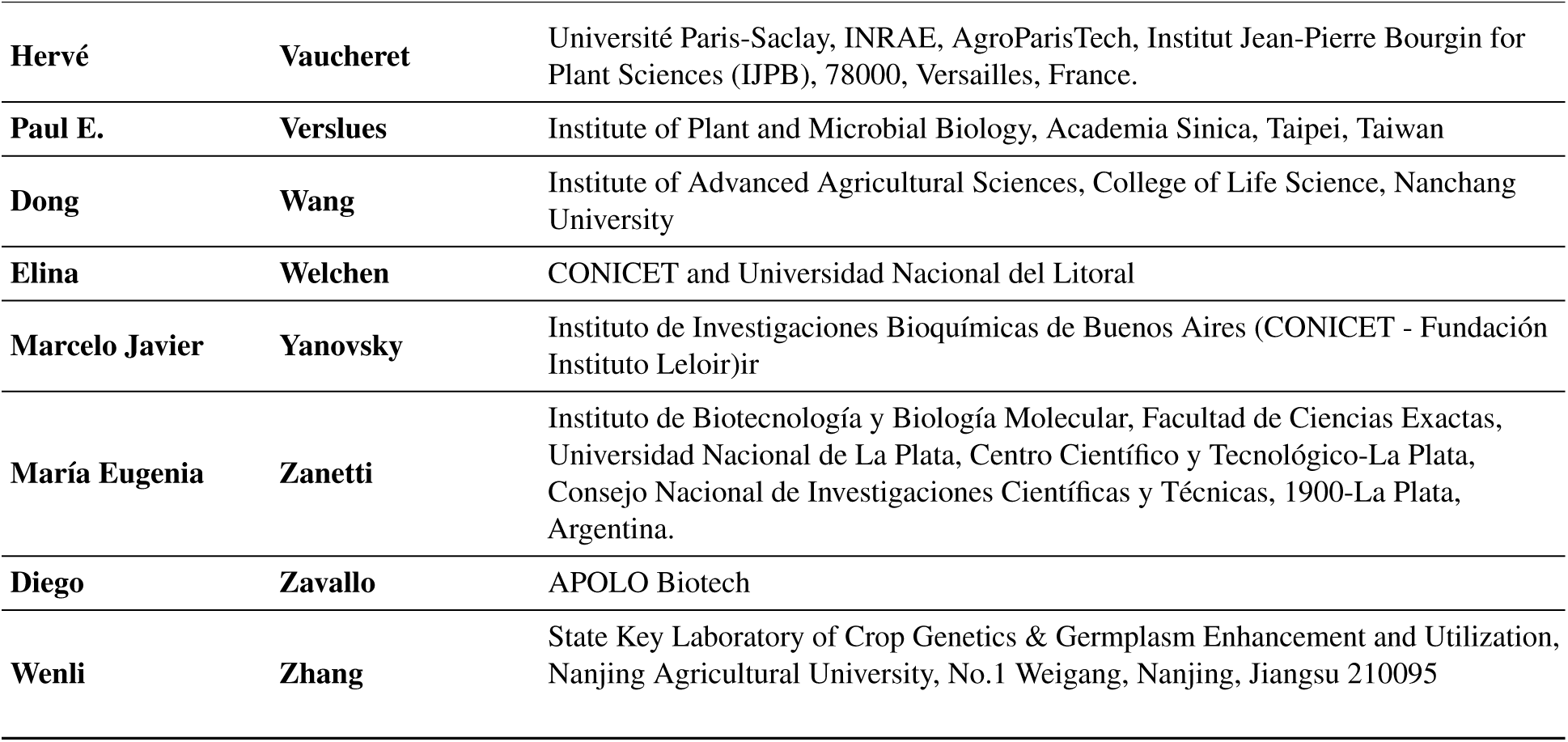
The MoBiPlant Consortium. All authors that comprised the confection and evaluation of MOBIPLANT are listed with their corresponding institution(s).

## B MOBIPLANT Examples

The following are 3 example questions drawn from **Expert MoBiPlant**.

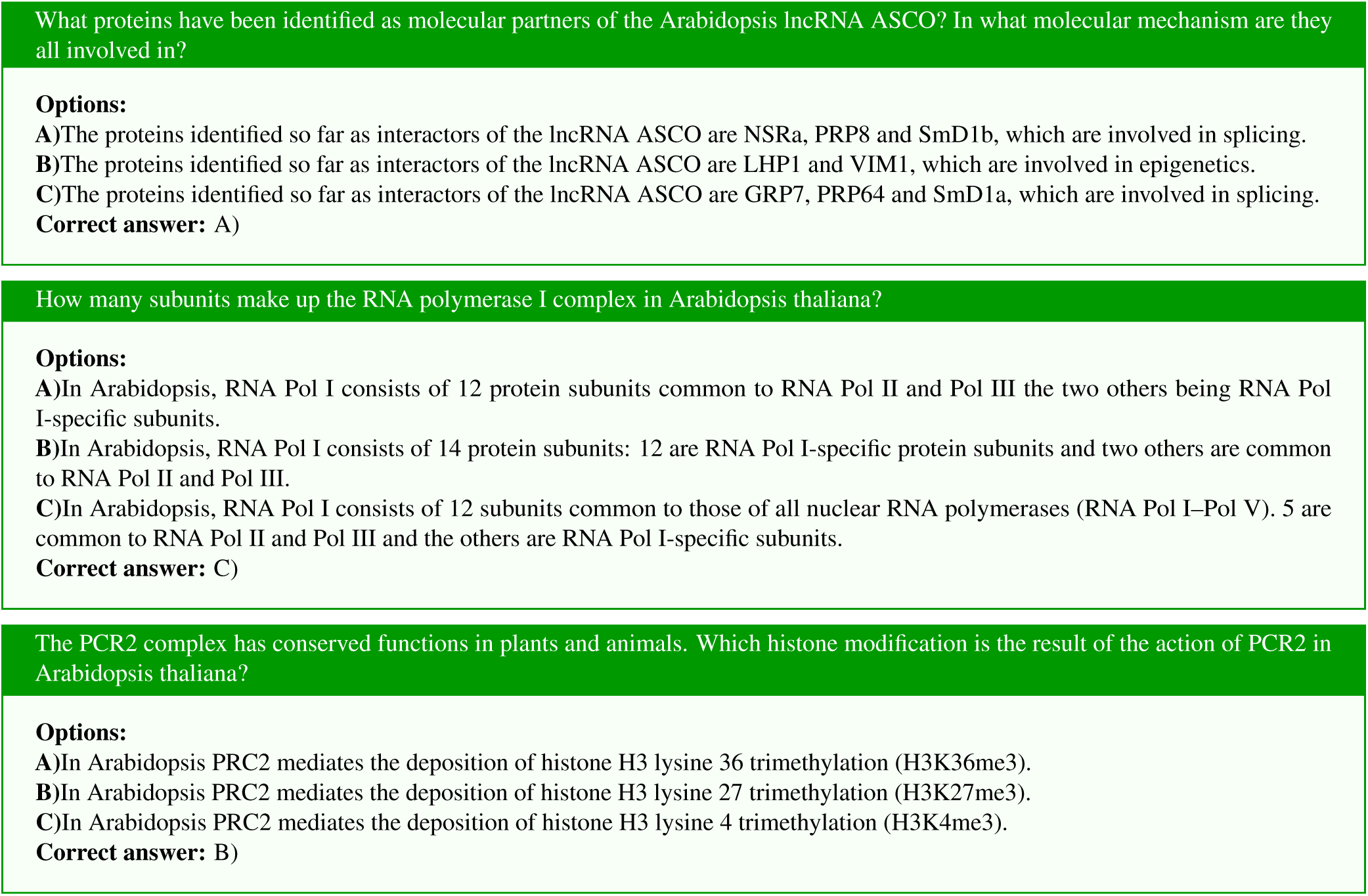

## C Synthetic MOBIPLANT Results

We report the evaluation results of Synthetic MoBiplant set in Supplementary Figure 1.

**Supplementary Figure 1.**
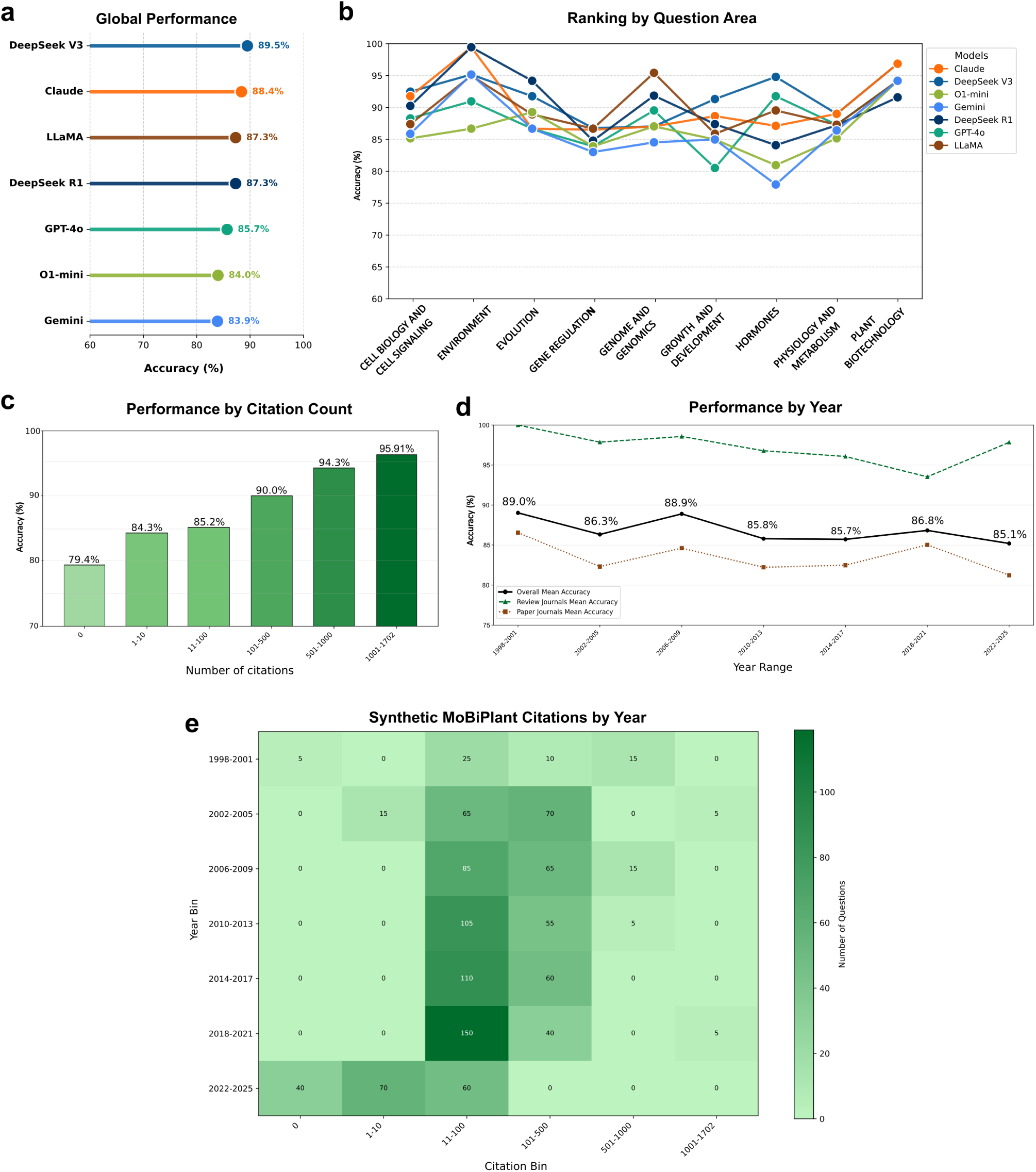
Synthetic MoBiPlant benchmarking results. **a)** The overall accuracy over the entire synthetic set. **b)** Fluctuation of model ranking across question areas. Each line and collection of dots represent the model accuracy in the area categories. **c)** Correlation between source’s amount of citations and mean model performance, with accuracy scores scaled to the 70–100 range. **d)** Accuracy plotted against publication year. The color intensity in the plot reflects the accuracy score for each data point. The source’s release year and citation count used in this analysis were extracted from the original work that motivated the research question.

## D Answer Distribution

Supplementary Table 2 shows the answer distribution of Gemini 1.5 Pro and DeepSeek-R1 in the option bias experiment.

**Supplementary Table 2.** Answer distributions for option bias experiment in MOBIPLANT. . For the original shuffles (no option bias), mean accuracy is reported across all three shuffles.

| Dataset & Model |  | A | B | C | Format Error | Answer Accuracy |
| --- | --- | --- | --- | --- | --- | --- |
| Option bias in A | GroundTruth | 565 | 0 | 0 | 0 | - |
|  | Gemini 1.5 Pro | 505 | 31 | 29 | 0 | 89.1 |
|  | DeepSeek R1 | 509 | 26 | 30 | 0 | 90.1 |
| Option bias in B | GroundTruth | 0 | 565 | 0 | 0 | - |
|  | Gemini 1.5 Pro | 120 | 397 | 32 | 16 | 71.9 |
|  | DeepSeek R1 | 58 | 470 | 37 | 0 | 82.9 |
| Option bias in C | GroundTruth | 0 | 0 | 565 | 0 | - |
|  | Gemini 1.5 Pro | 140 | 55 | 356 | 14 | 64.3 |
|  | DeepSeek R1 | 61 | 44 | 460 | 0 | 81.0 |
| Original Gemini 1.5 Pro | First shuffle | 262 | 163 | 140 | 0 | 76.8 |
|  | Second shuffle | 256 | 164 | 145 | 0 |  |
|  | Third shuffle | 264 | 154 | 147 | 0 |  |
| Original DeepSeek R1 | First shuffle | 225 | 168 | 168 | 4 | 86.1 |
|  | Second shuffle | 208 | 168 | 174 | 15 |  |
|  | Third shuffle | 226 | 155 | 174 | 10 |  |

## E Failure Modes

In Supplementary Table 3 we report additional examples of failure modes encountered on the models evaluated on MOBIPLANT. We extend the definition of the found set of failure modes on the following list:

- **Cross-domain bias**: Apply animal or human biology mechanisms to plants incorrectly.
- **Species confusion**: Mix up information across different plant species.
- **Acronym misinterpretation**: Expand plant-specific abbreviations into wrong or non-existent gene names.
- **Terminology gaps**: Misunderstand specialized plant physiology terms (e.g. thermomorphogenesis).
- **Outdated knowledge**: Lack recent literature on emerging plant research areas.
- **Hallucinated pathways**: Combine unrelated defense or signaling pathways or involve incorrect genes in a given molecular mechanism.
- **Environmental misunderstanding**: Struggle with field conditions, temperature stress, and abiotic factors.

**Supplementary Table 3.**
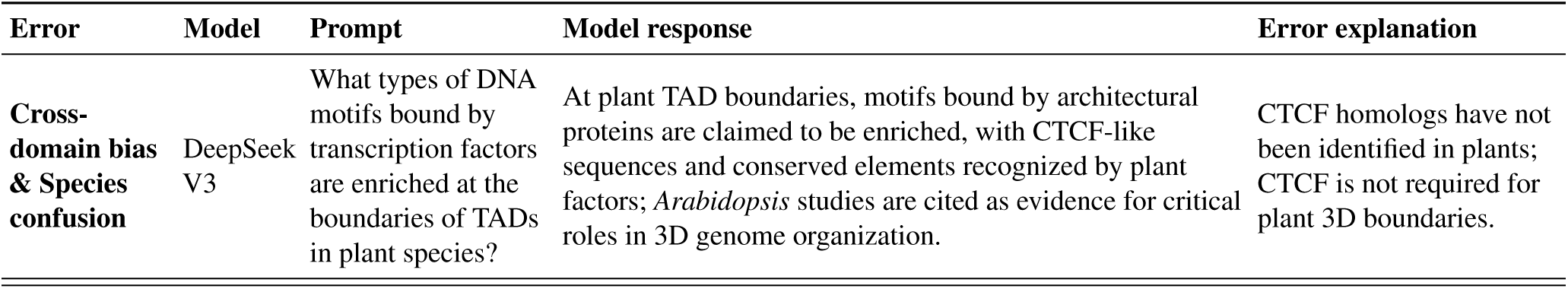
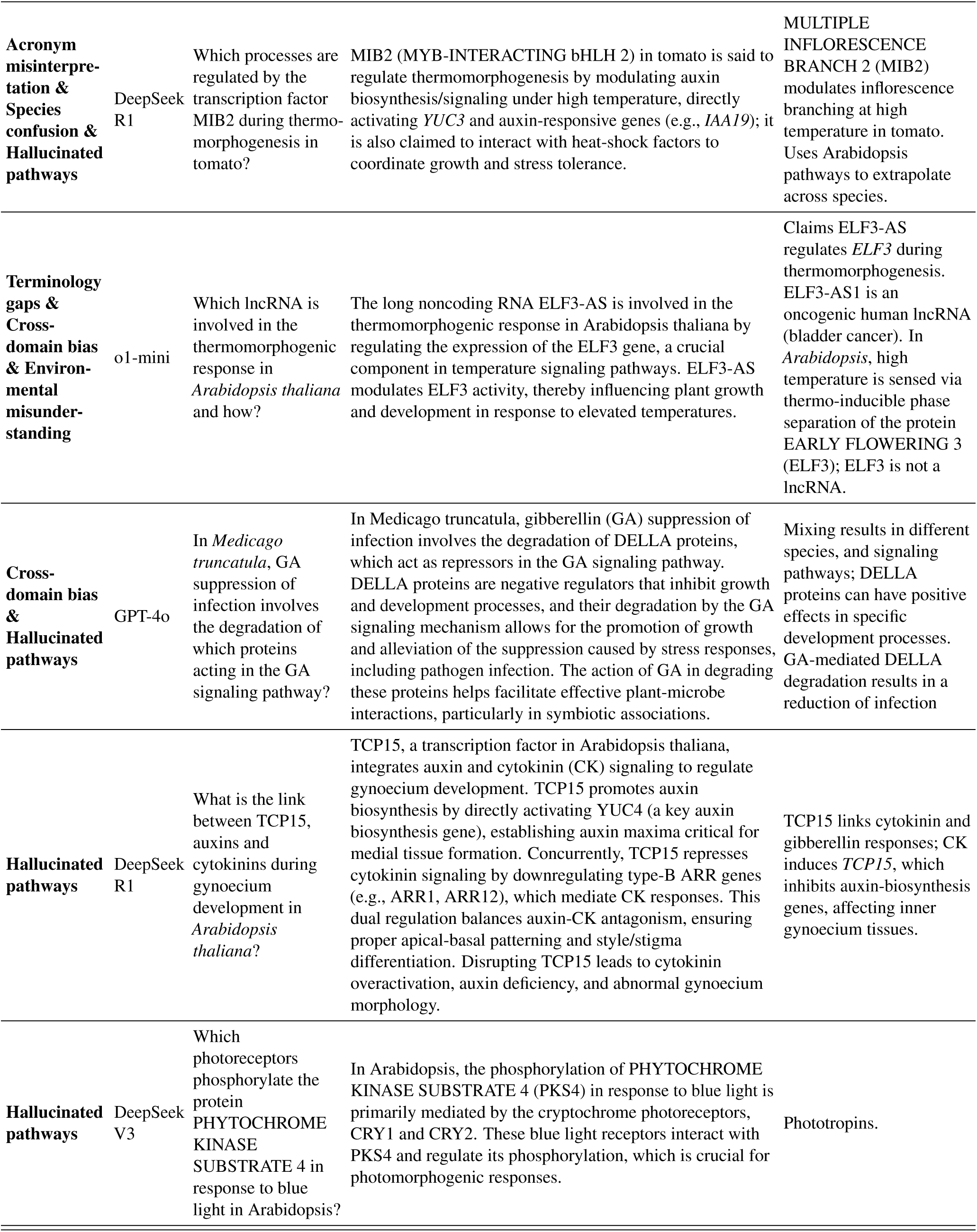
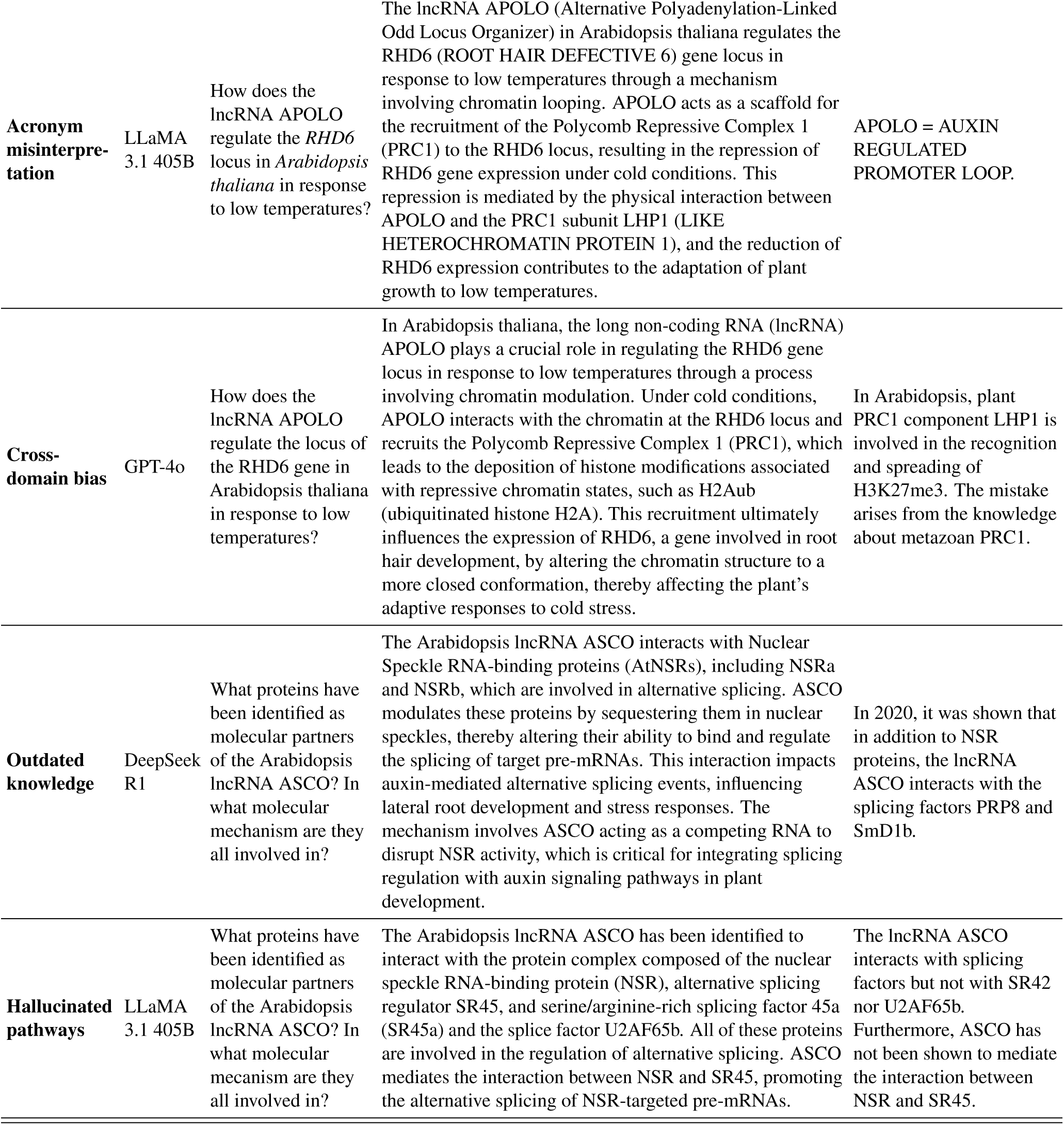
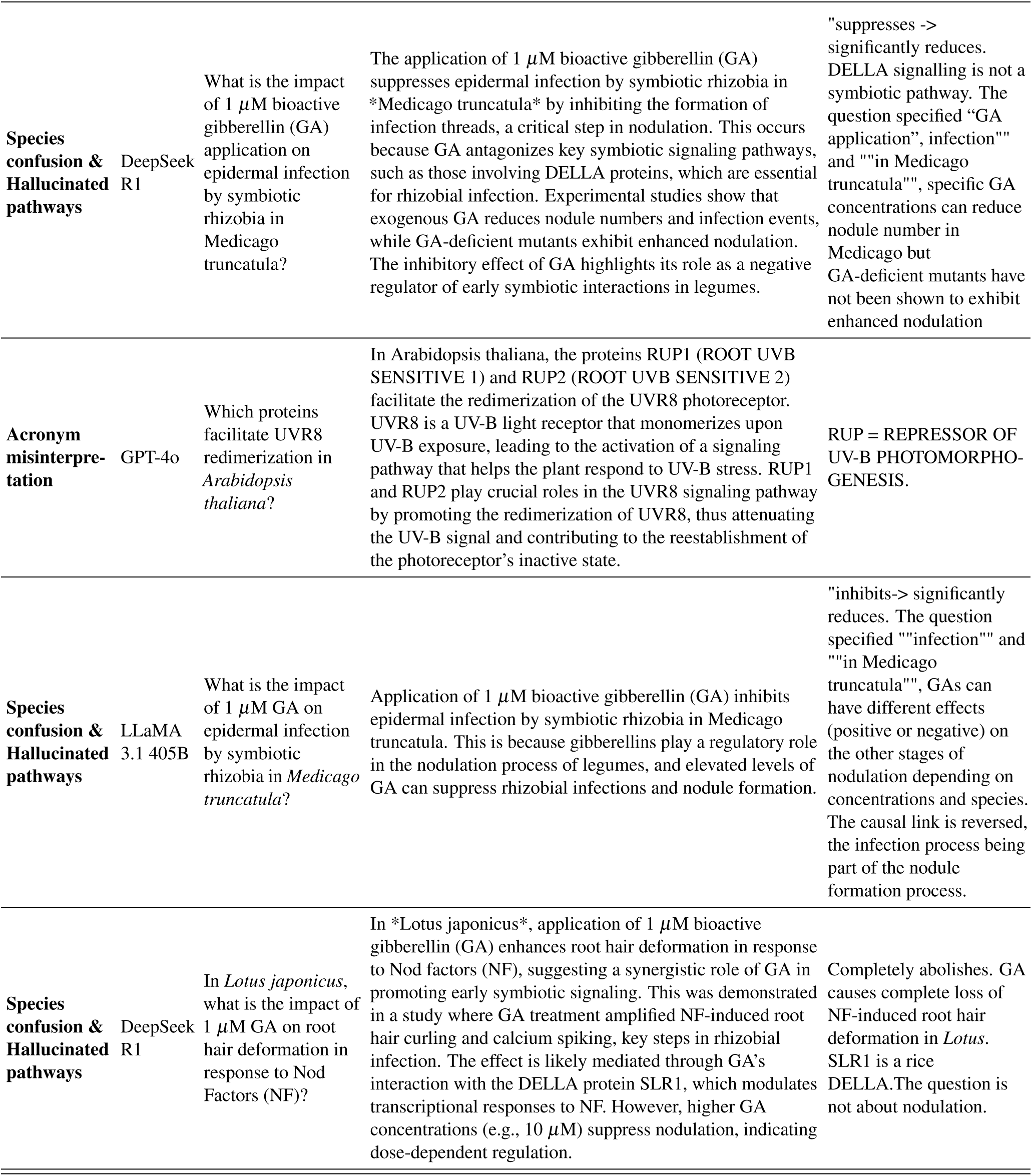
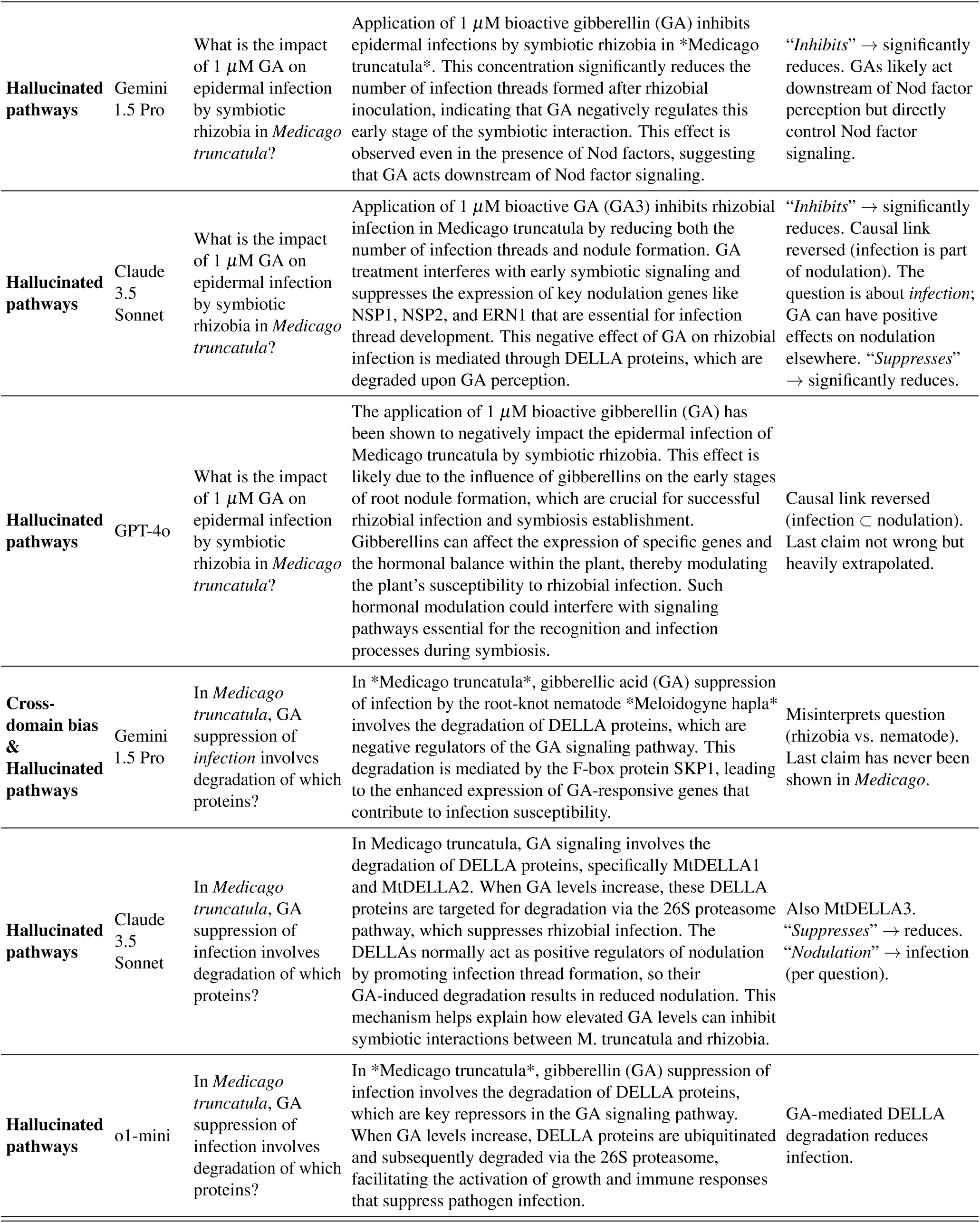
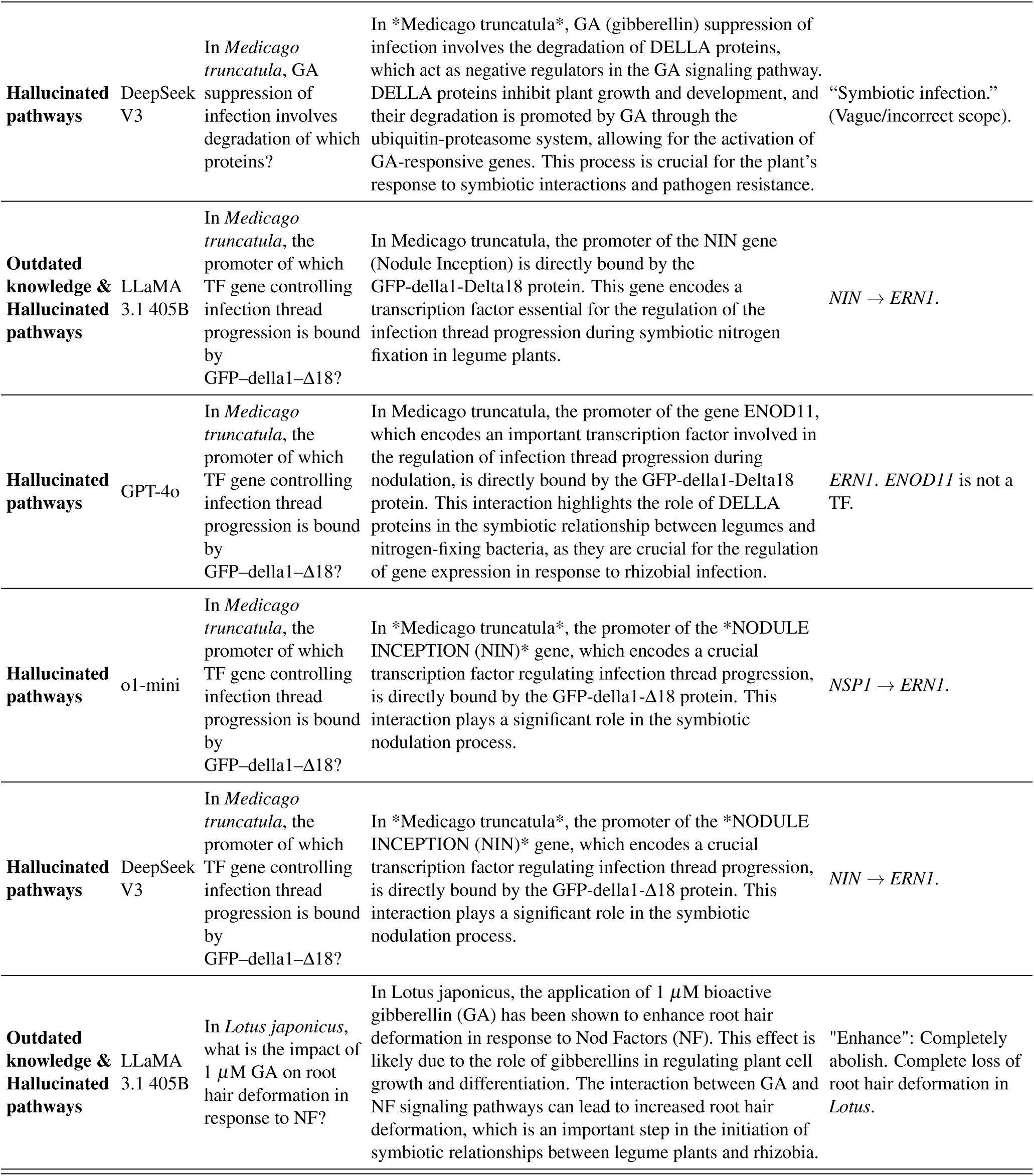
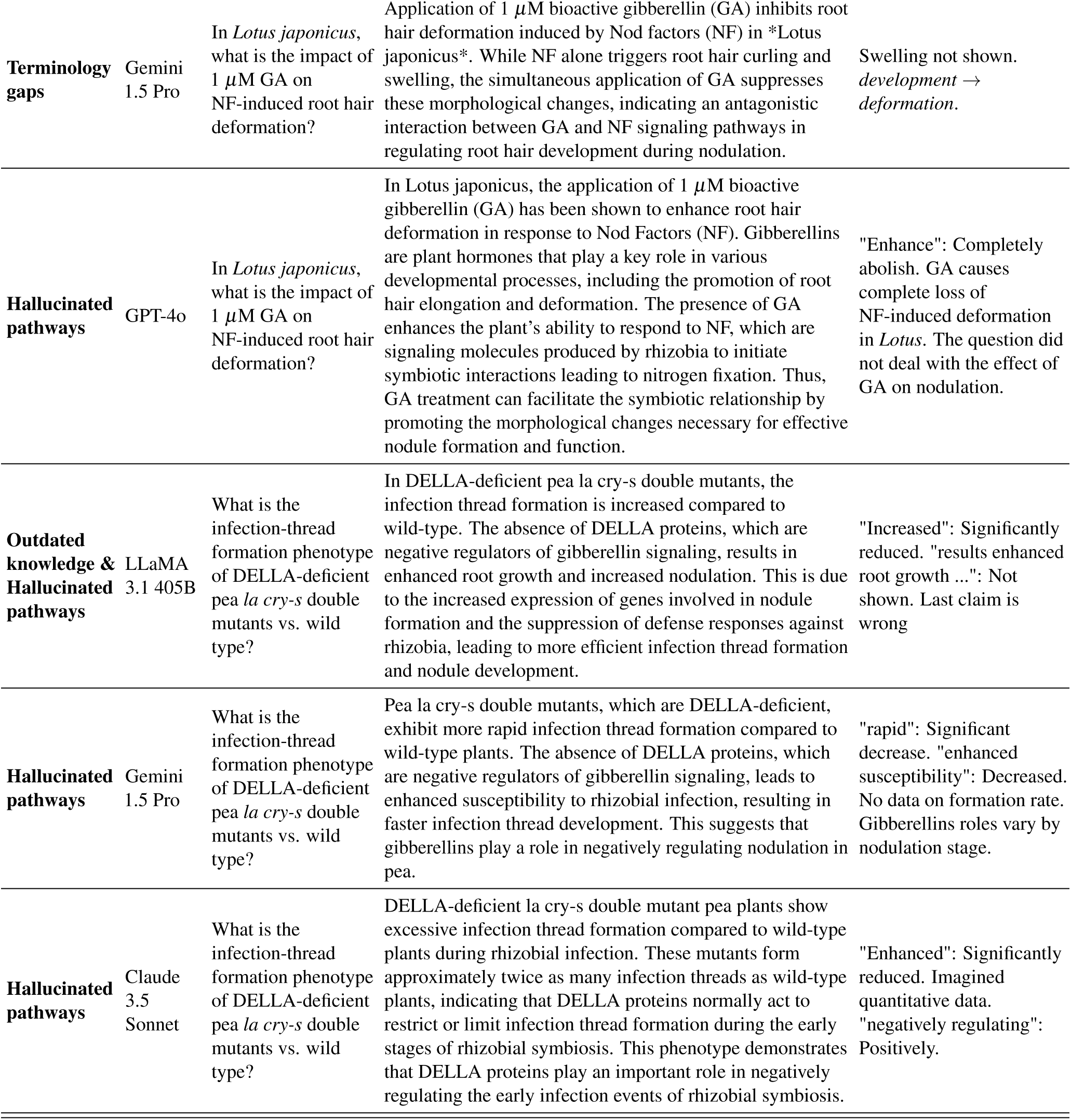
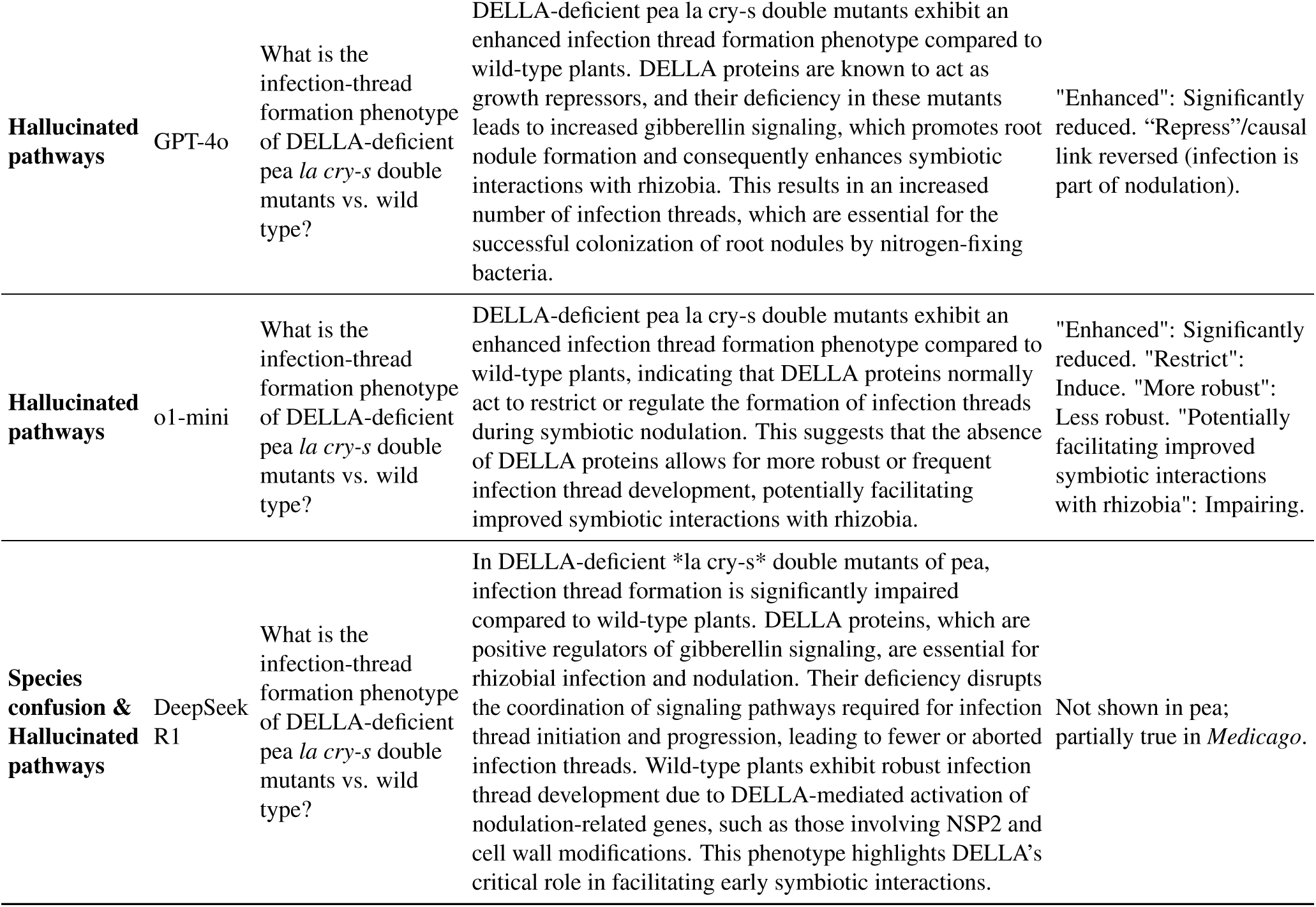
Additional failure-mode examples from the open-ended assessment of Expert MoBiPlant.

**Supplementary Table 4.**
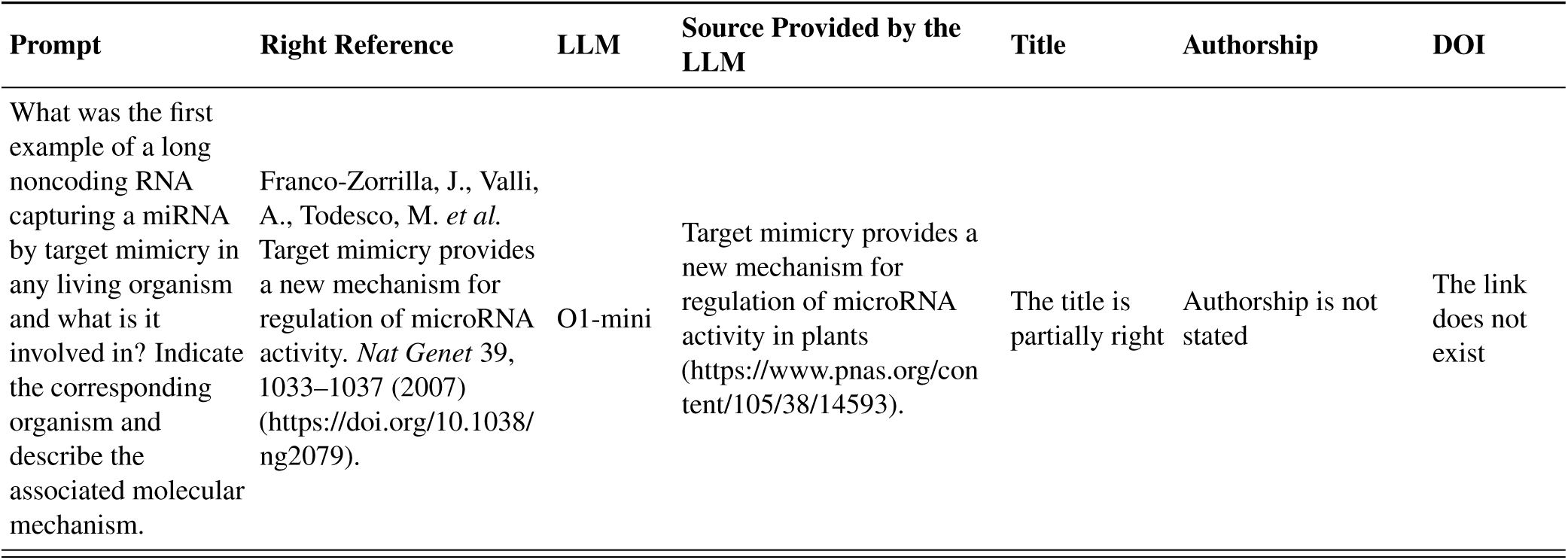
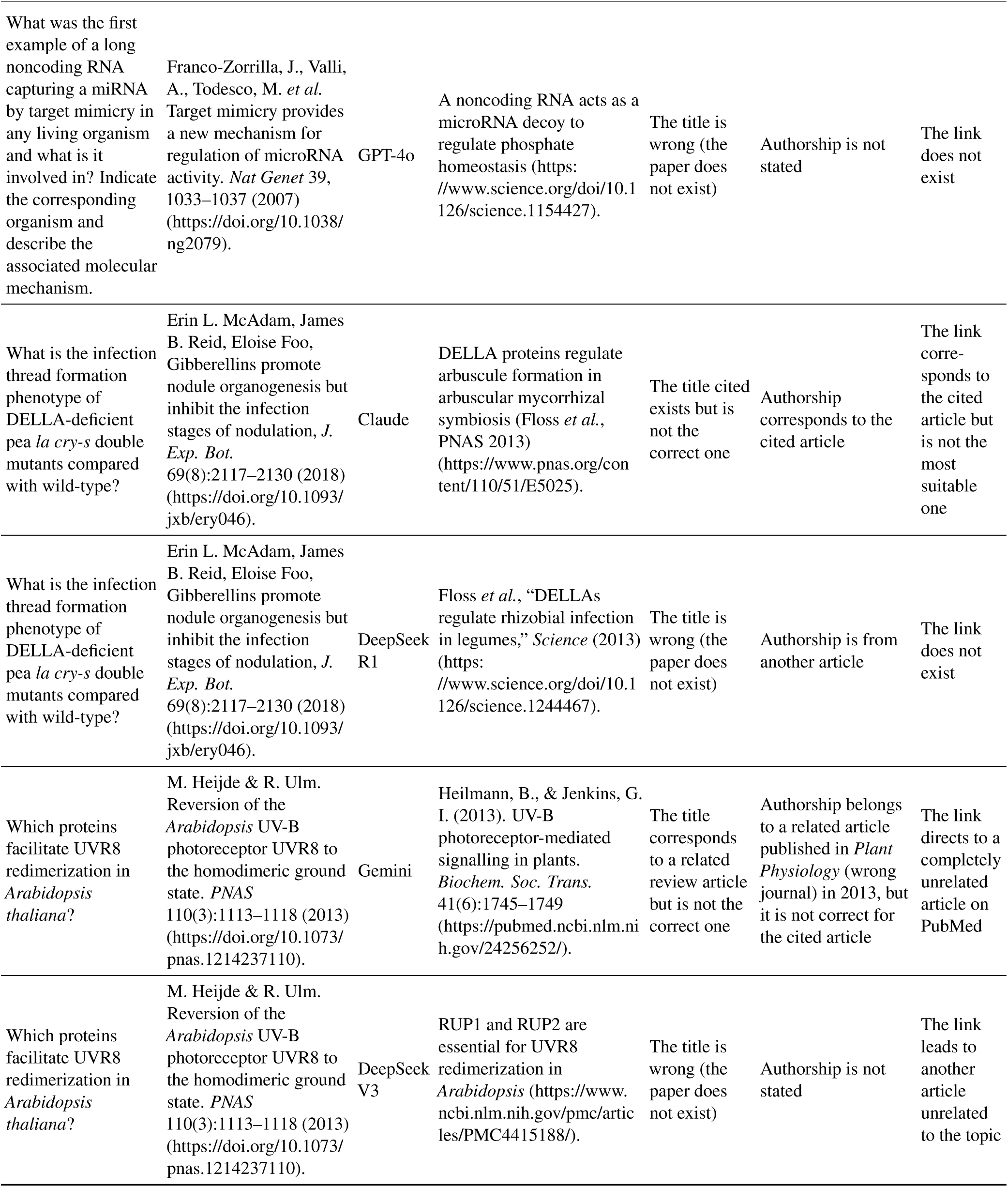
Examples of common failure modes sighted on the open-ended assessment of Expert MoBiPlant for the references indicated by LLMs to support their statements.

| Prompt | Right Reference | LLM | Source Provided by the LLM | Title | Authorship | DOI |
| --- | --- | --- | --- | --- | --- | --- |
| What was the first example of a long noncoding RNA capturing a miRNA by target mimicry in any living organism and what is it involved in? Indicate the corresponding organism and describe the associated molecular mechanism. | Franco-Zorrilla, J., Valli, A., Todesco, M. *et al.* Target mimicry provides a new mechanism for regulation of microRNA activity. *Nat Genet* 39, 1033–1037 (2007) (https://doi.org/10.1038/ng2079). | O1-mini | Target mimicry provides a new mechanism for regulation of microRNA activity in plants (https://www.pnas.org/content/105/38/14593). | The title is partially right | Authorship is not stated | The link does not exist |
| What was the first example of a long noncoding RNA capturing a miRNA by target mimicry in any living organism and what is it involved in? Indicate the corresponding organism and describe the associated molecular mechanism. | Franco-Zorrilla, J., Valli, A., Todesco, M. *et al.* Target mimicry provides a new mechanism for regulation of microRNA activity. *Nat Genet* 39, 1033–1037 (2007) (https://doi.org/10.1038/ng2079). | GPT-4o | A noncoding RNA acts as a microRNA decoy to regulate phosphate homeostasis (https://www.science.org/doi/10.1126/science.1154427). | The title is wrong (the paper does not exist) | Authorship is not stated | The link does not exist |
| What is the infection thread formation phenotype of DELLA-deficient pea *la cry-s* double mutants compared with wild-type? | Erin L. McAdam, James B. Reid, Eloise Foo, Gibberellins promote nodule organogenesis but inhibit the infection stages of nodulation, *J. Exp. Bot.* 69(8):2117–2130 (2018) (https://doi.org/10.1093/jxb/ery046). | Claude | DELLA proteins regulate arbuscule formation in arbuscular mycorrhizal symbiosis (Floss *et al.*, PNAS 2013) (https://www.pnas.org/content/110/51/E5025). | The title cited exists but is not the correct one | Authorship corresponds to the cited article | The link corresponds to the cited article but is not the most suitable one |
| What is the infection thread formation phenotype of DELLA-deficient pea *la cry-s* double mutants compared with wild-type? | Erin L. McAdam, James B. Reid, Eloise Foo, Gibberellins promote nodule organogenesis but inhibit the infection stages of nodulation, *J. Exp. Bot.* 69(8):2117–2130 (2018) (https://doi.org/10.1093/jxb/ery046). | DeepSeek R1 | Floss *et al.*, “DELLAs regulate rhizobial infection in legumes,” *Science* (2013) (https://www.science.org/doi/10.1126/science.1244467). | The title is wrong (the paper does not exist) | Authorship is from another article | The link does not exist |
| Which proteins facilitate UVR8 redimerization in *Arabidopsis thaliana*? | M. Heijde & R. Ulm. Reversion of the *Arabidopsis* UV-B photoreceptor UVR8 to the homodimeric ground state. *PNAS* 110(3):1113–1118 (2013) (https://doi.org/10.1073/pnas.1214237110). | Gemini | Heilmann, B., & Jenkins, G. I. (2013). UV-B photoreceptor-mediated signalling in plants. *Biochem. Soc. Trans.* 41(6):1745–1749 (https://pubmed.ncbi.nlm.nih.gov/24256252/). | The title corresponds to a related review article but is not the correct one | Authorship belongs to a related article published in *Plant Physiology* (wrong journal) in 2013, but it is not correct for the cited article | The link directs to a completely unrelated article on PubMed |
| Which proteins facilitate UVR8 redimerization in *Arabidopsis thaliana*? | M. Heijde & R. Ulm. Reversion of the *Arabidopsis* UV-B photoreceptor UVR8 to the homodimeric ground state. *PNAS* 110(3):1113–1118 (2013) (https://doi.org/10.1073/pnas.1214237110). | DeepSeek V3 | RUP1 and RUP2 are essential for UVR8 redimerization in *Arabidopsis* (https://www.ncbi.nlm.nih.gov/pmc/articles/PMC4415188/). | The title is wrong (the paper does not exist) | Authorship is not stated | The link leads to another article unrelated to the topic |

## F Agentic LLMs and DNA models on MOBIPLANT

We show Claude models and ChatGPT models along ChatNT performances on MOBIPLANT in Figure 2. For the OpenAI family, GPT-5.2 achieves 90.9 ± 1.2%, a substantially larger gain over GPT-4o (81.0 ± 2.1%). ChatNT achieves a substantially lower overall accuracy, primarily due to the reduced size of its language encoder (inherited from Vicuna 7B) compared to the closed models, which limits its reasoning capabilities.

**Supplementary Figure 2.**
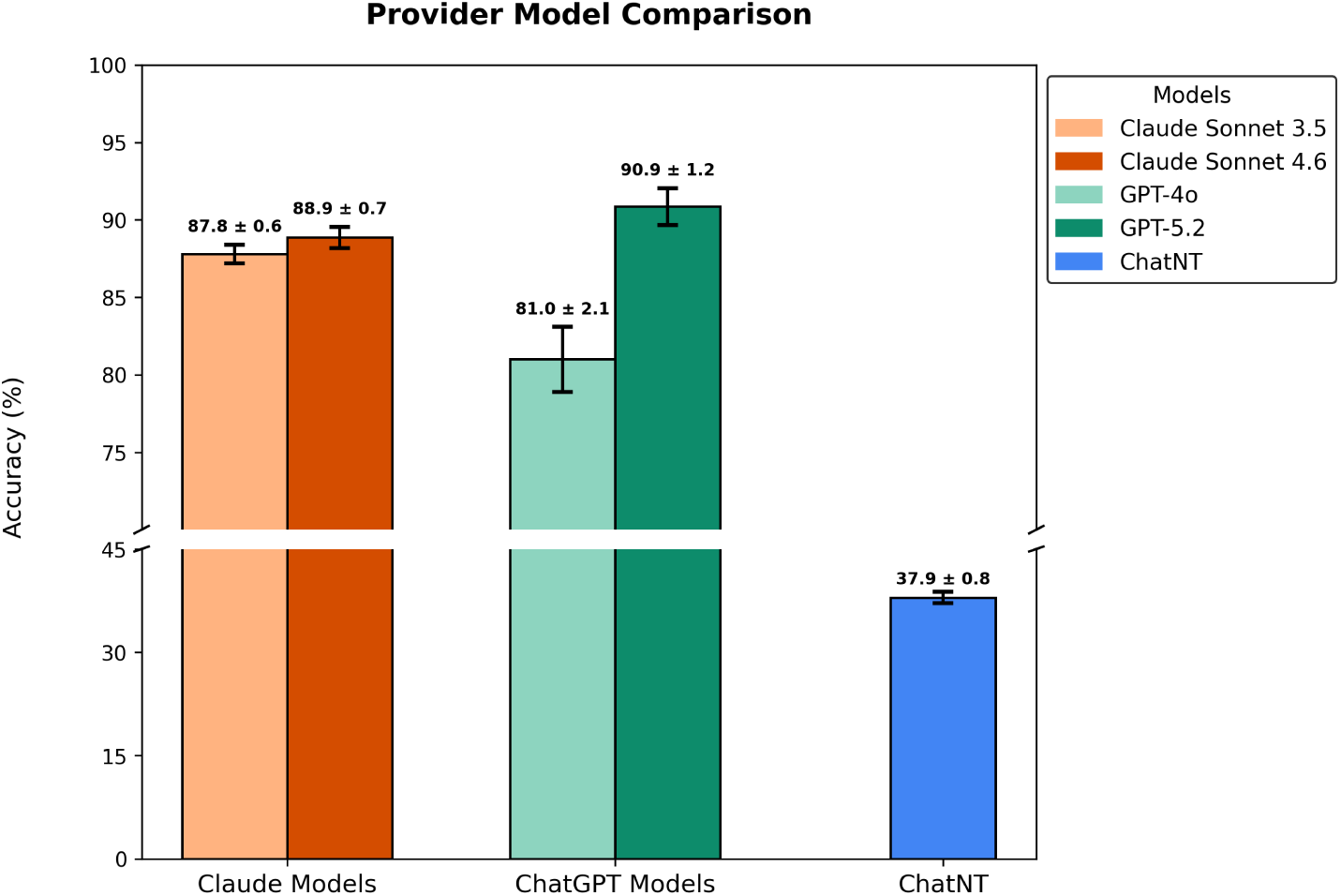
Performance comparison of successive model generations on Expert MoBiPlant. Accuracy (%) of two successive model versions with web access for each of two providers on the Expert MoBiPlant set, along ChatNT. Error bars represent standard deviations across three independent repetitions with shuffled answer option order.

## G Plant DNA Queries

**Supplementary Figure 3.**
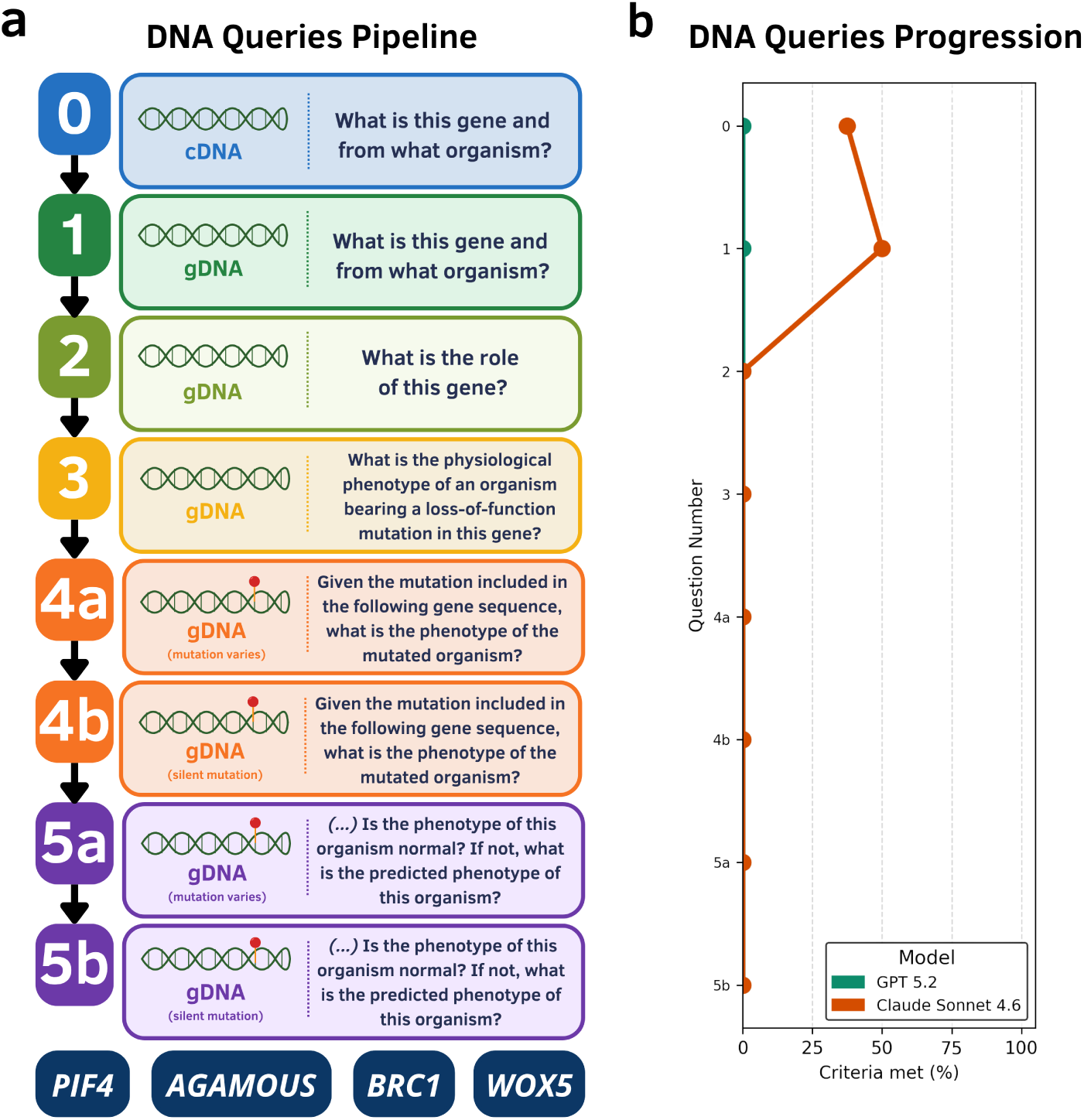
Stepwise evaluation of large language model performance on DNA-based gene reasoning tasks. **a)** Schematic of the DNA Queries Pipeline, comprising eight sequential prompts of increasing biological complexity. Steps 0 and 1 ask the model to identify the gene and its source organism from a cDNA and a gDNA sequence, respectively. Step 2 queries the functional role of the gene given its gDNA sequence. Step 3 probes the ability to infer the physiological phenotype resulting from a loss-of-function mutation based on gDNA sequence identity. Steps 4a and 4b provide a gDNA sequence carrying either a deletion/substitution/addition (4a) or silent point mutation (4b) and ask the model to predict the phenotype of the mutant organism. Steps 5a and 5b present a gDNA sequence harbouring a deletion/substitution/addition (5a) or silent mutation (5b) without explicitly stating whether the sequence is mutated, requiring the model to recognize the presence and nature of the mutation and infer whether the organism’s phenotype is expected to deviate from wild type. Each query in the pipeline was applied to four plant genes representative of distinct regulatory functions: PIF4, AGAMOUS, BRC1, and WOX5. **b)** Performance of GPT 5.2 (teal) and Claude Sonnet 4.6 (orange) at each step of the pipeline, expressed as the percentage of evaluation criteria scored (averaged across the four genes). ChatNT was not included in this evaluation, as it returned outputs unrelated to the queries when prompted in this open-ended format, consistent with its fine-tuning on constrained DNA classification and regression templates.

## H MCQ Collection Guidelines

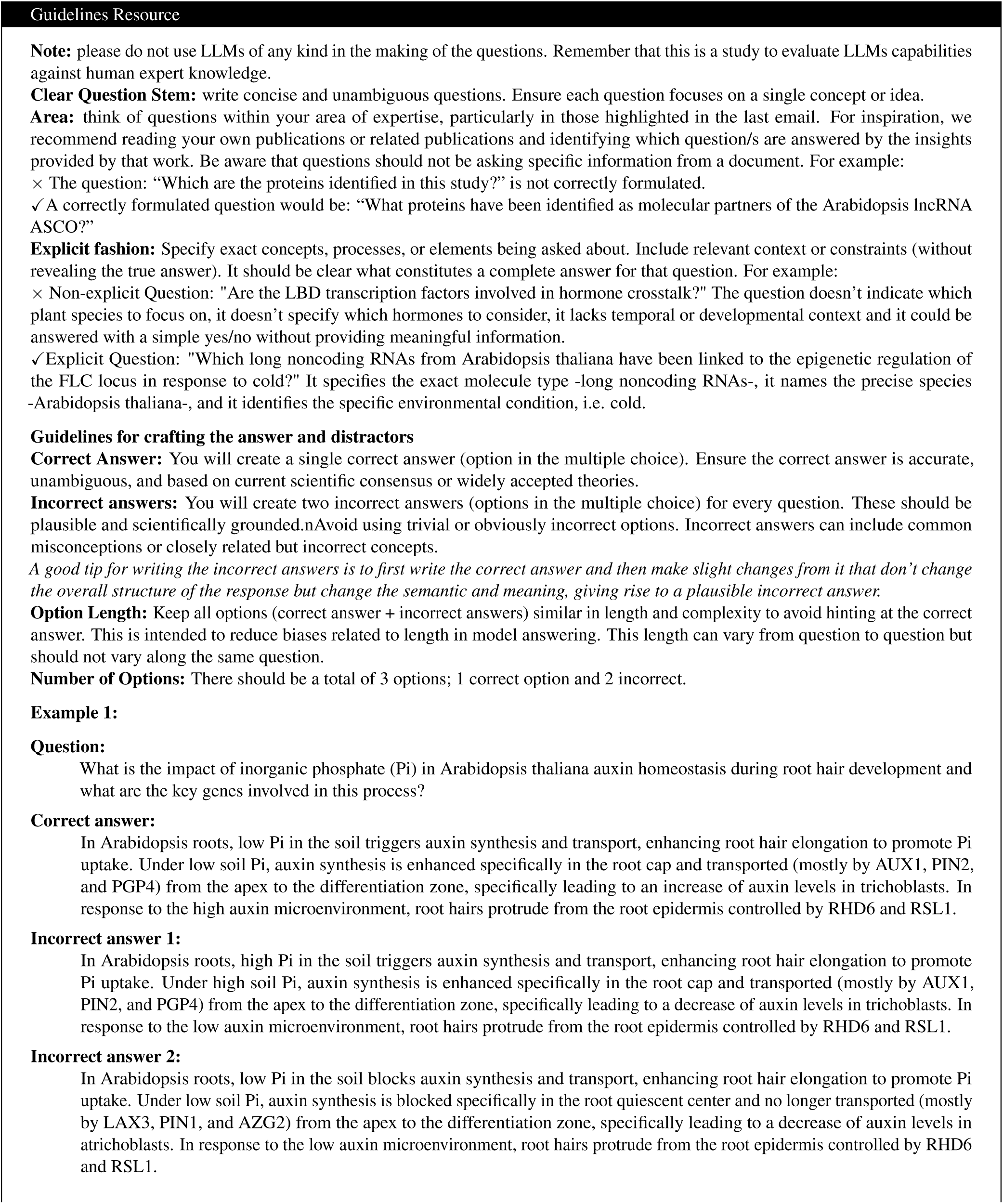
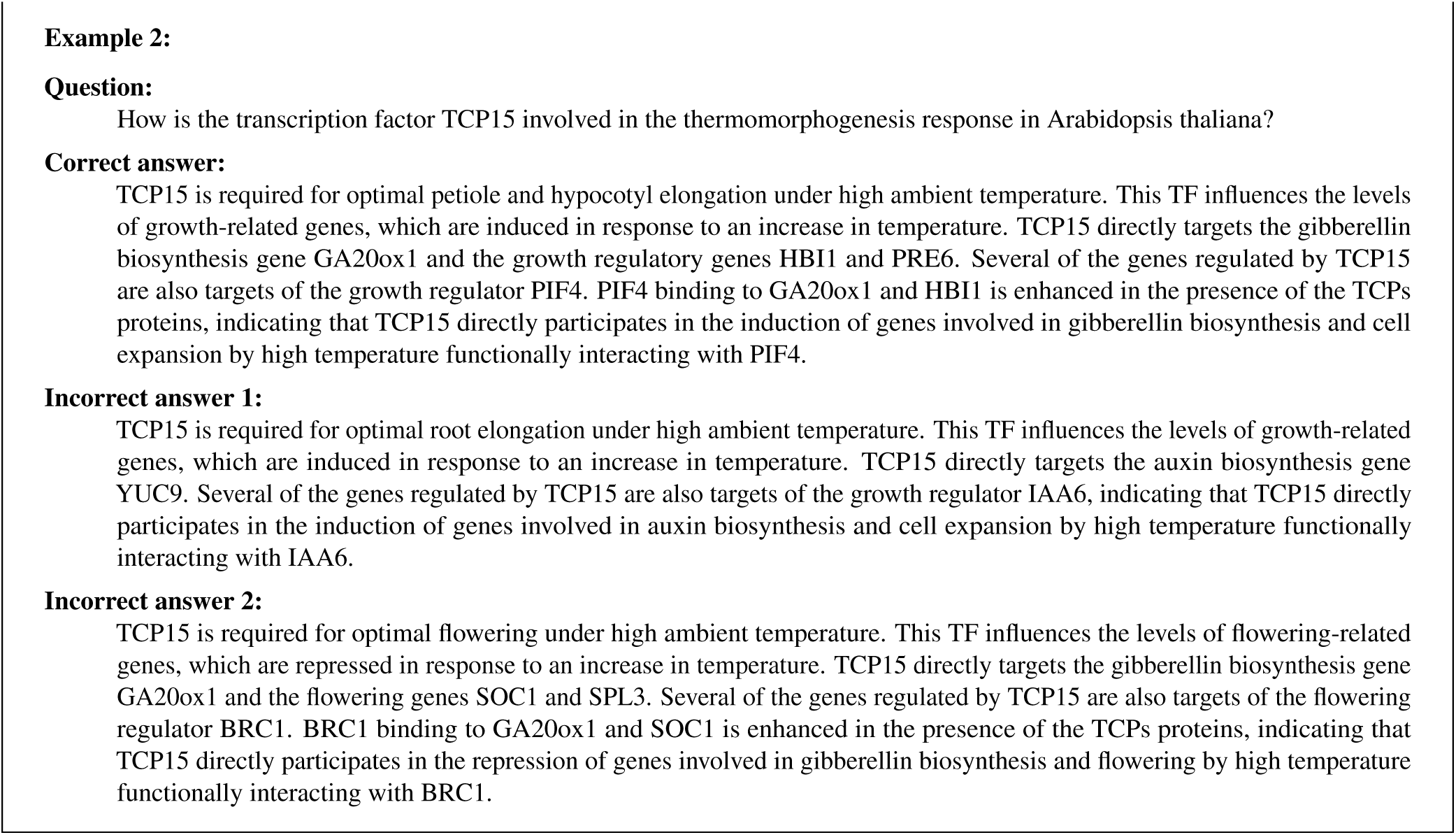

## I Prompts

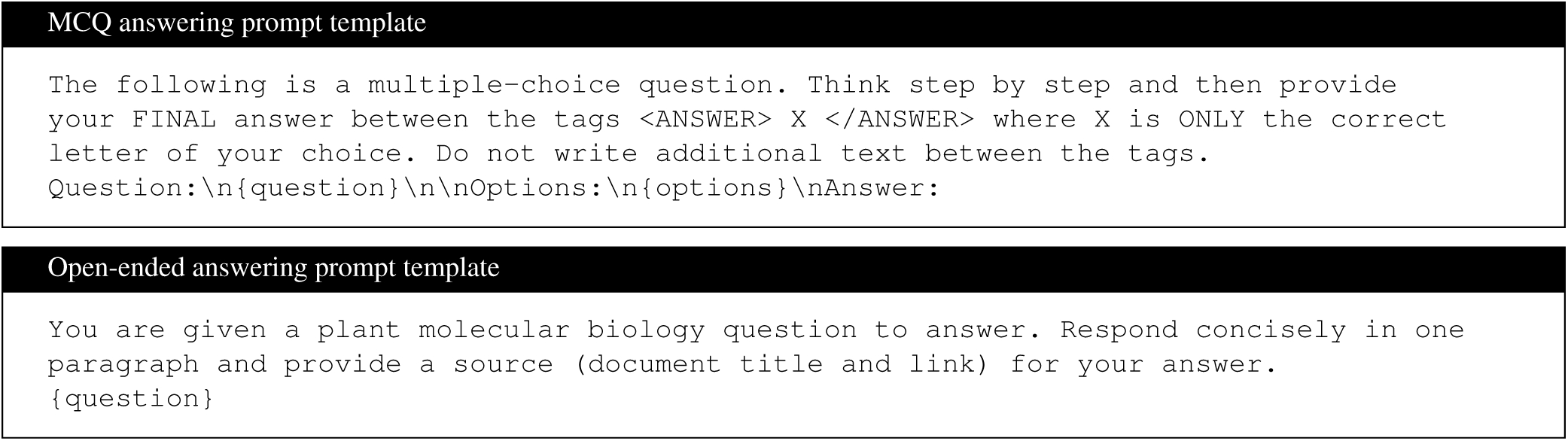

The following is the prompt instruction template used to generate Synthetic MoBiPlant.

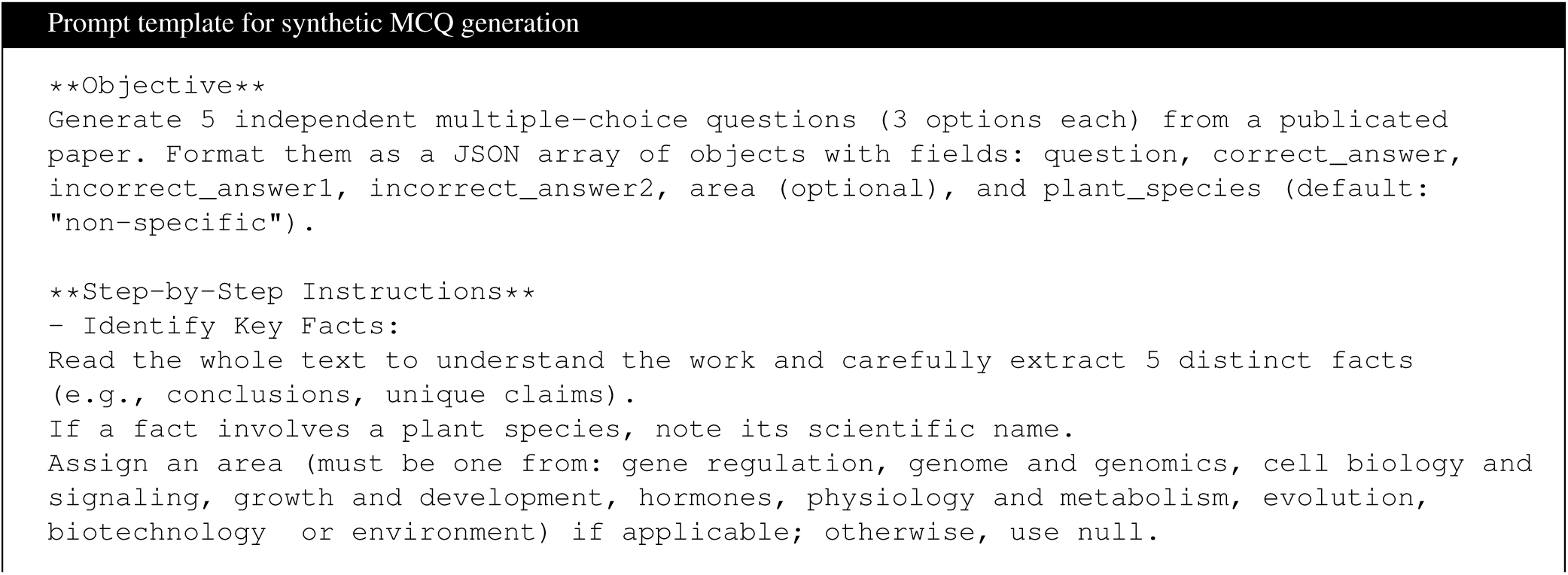
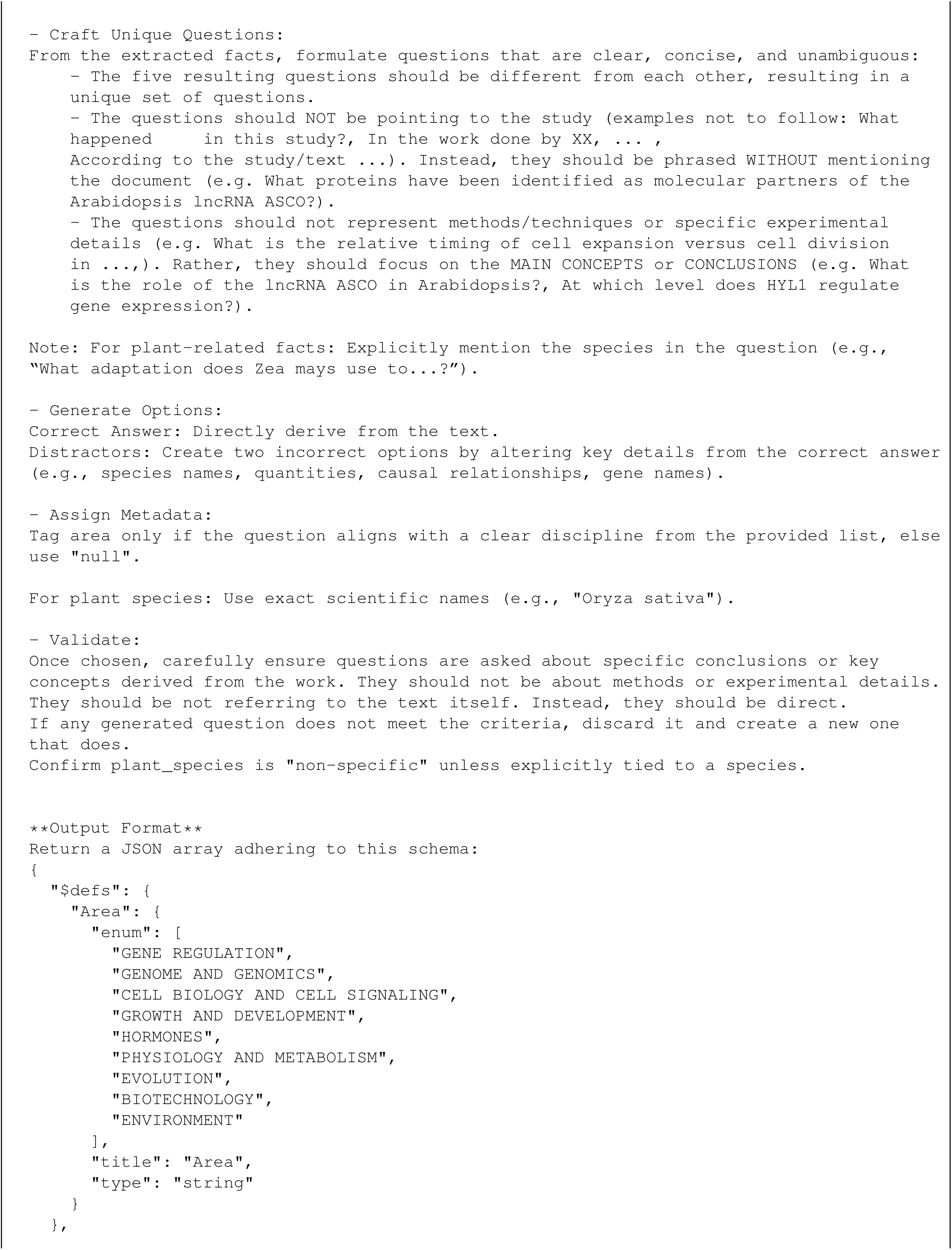
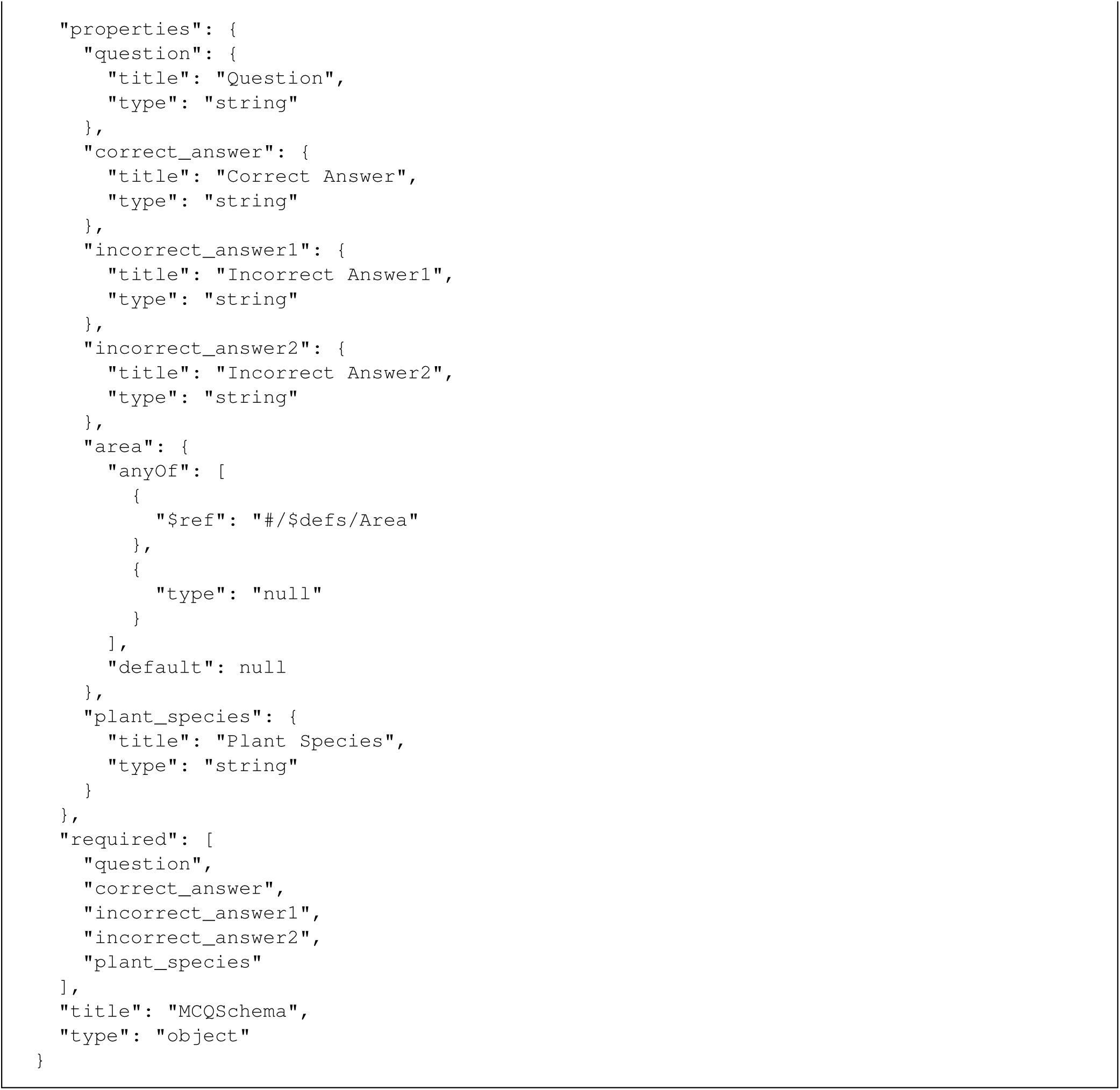

The following is the prompt instruction template used to evaluate the Plant DNA Queries using GPT 5.2.

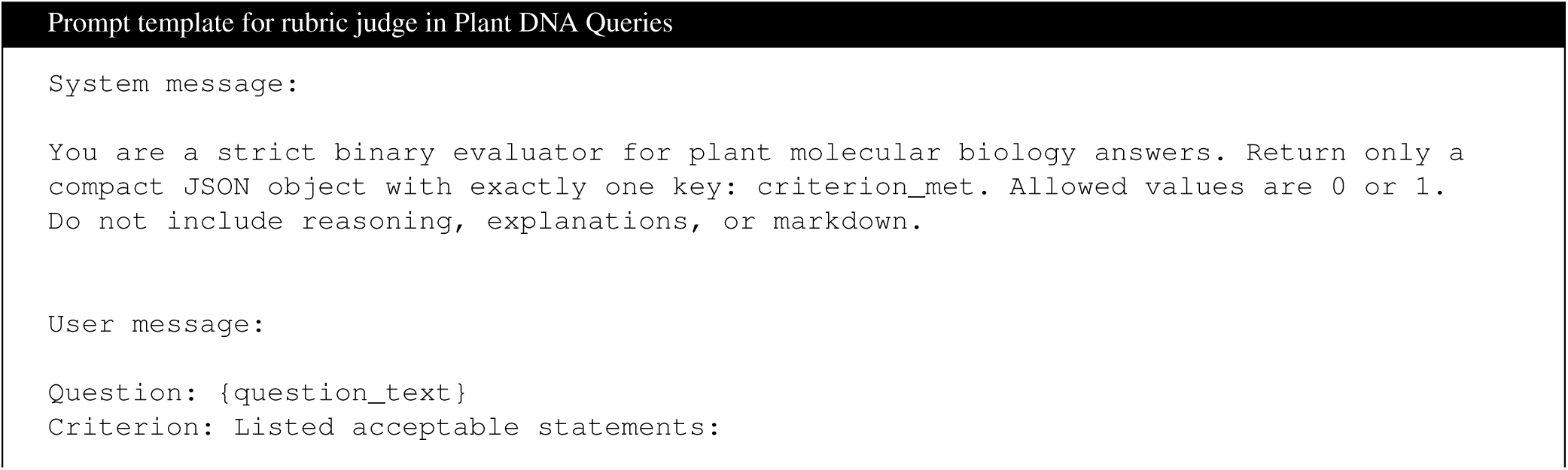
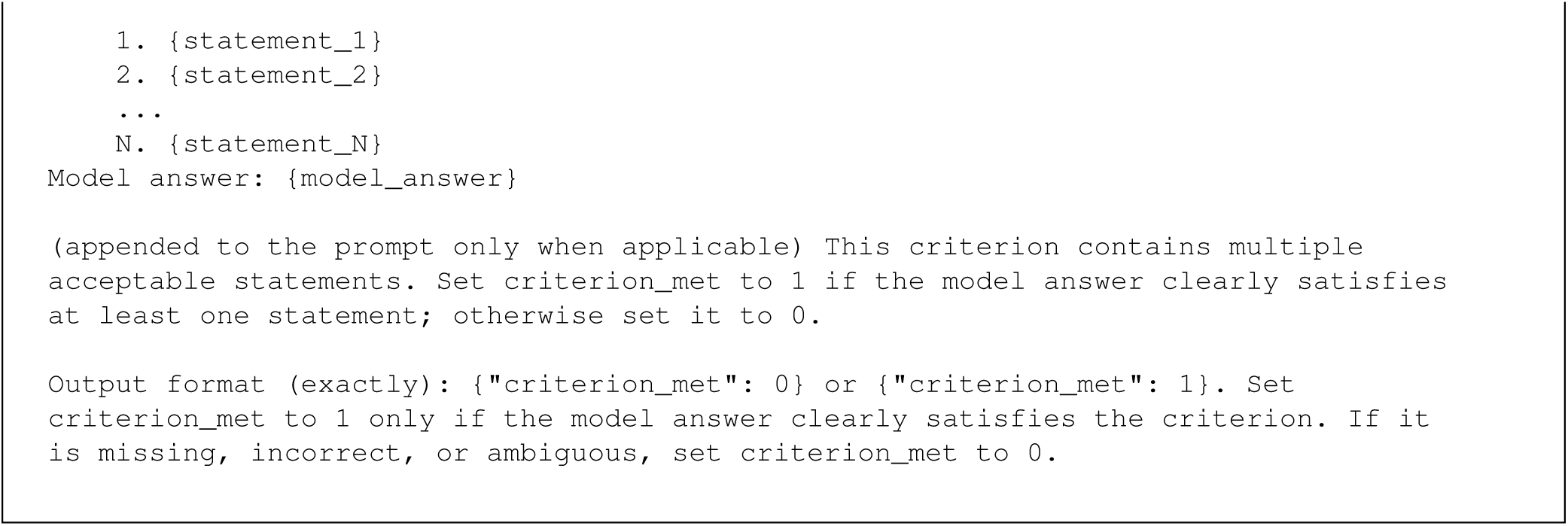

## J Plant DNA Queries Responses & Rubric

We show the model responses to our Plant DNA Queries dataset in Supplementary Table 5. On Supplementary Table 7, we detail the rubrics used for scoring open ended responses to these questions. Supplementary Tables 5 and 7 are appended as separate xlsx files.

## K Journal Distribution

We report the journal distribution of the Expert MoBiPlant set in Supplementary Table 6 and the models’ performance on the journal performance experiment in Supplementary Table 8, along the journal distribution.

**Supplementary Table 6.**
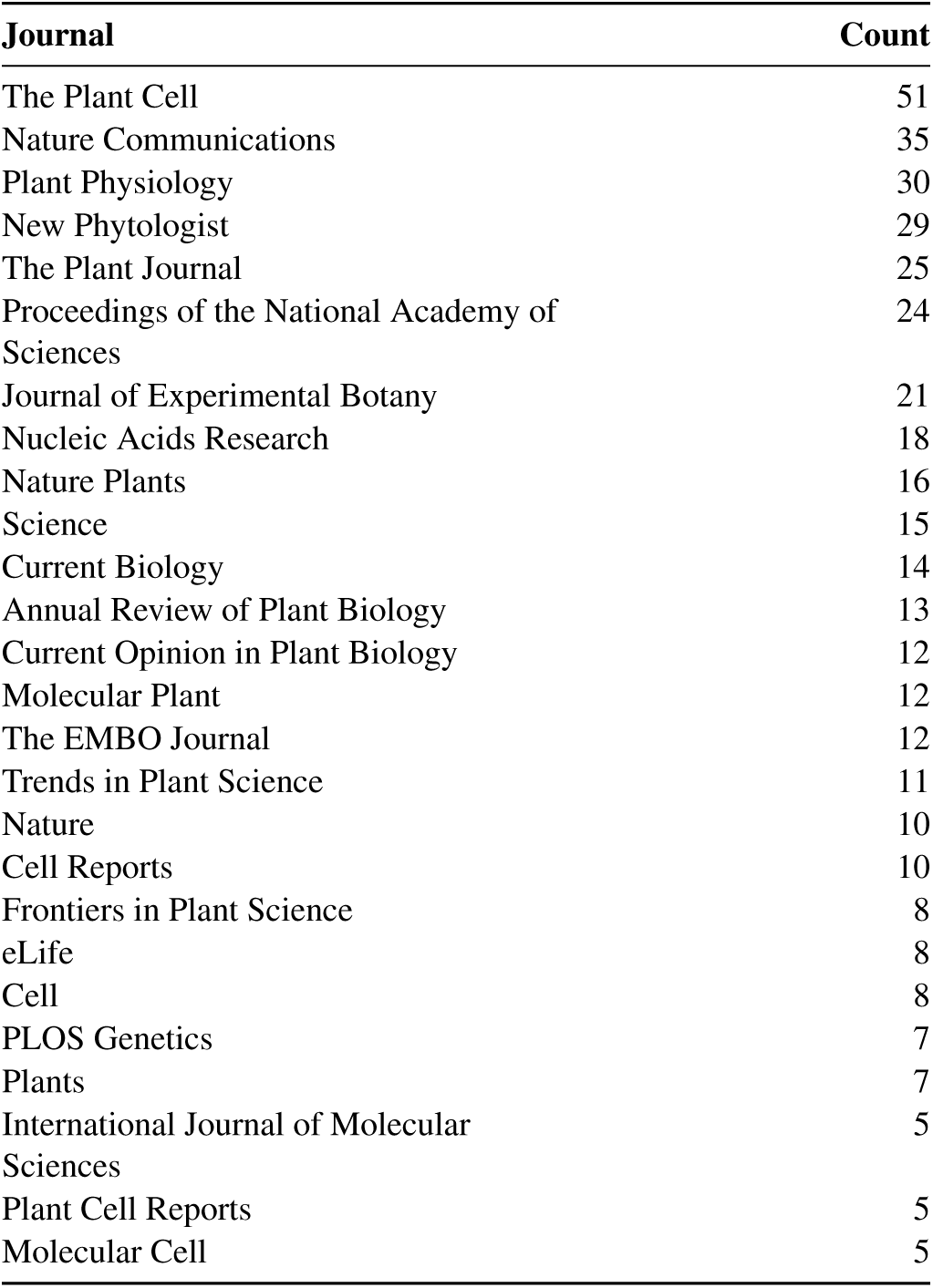
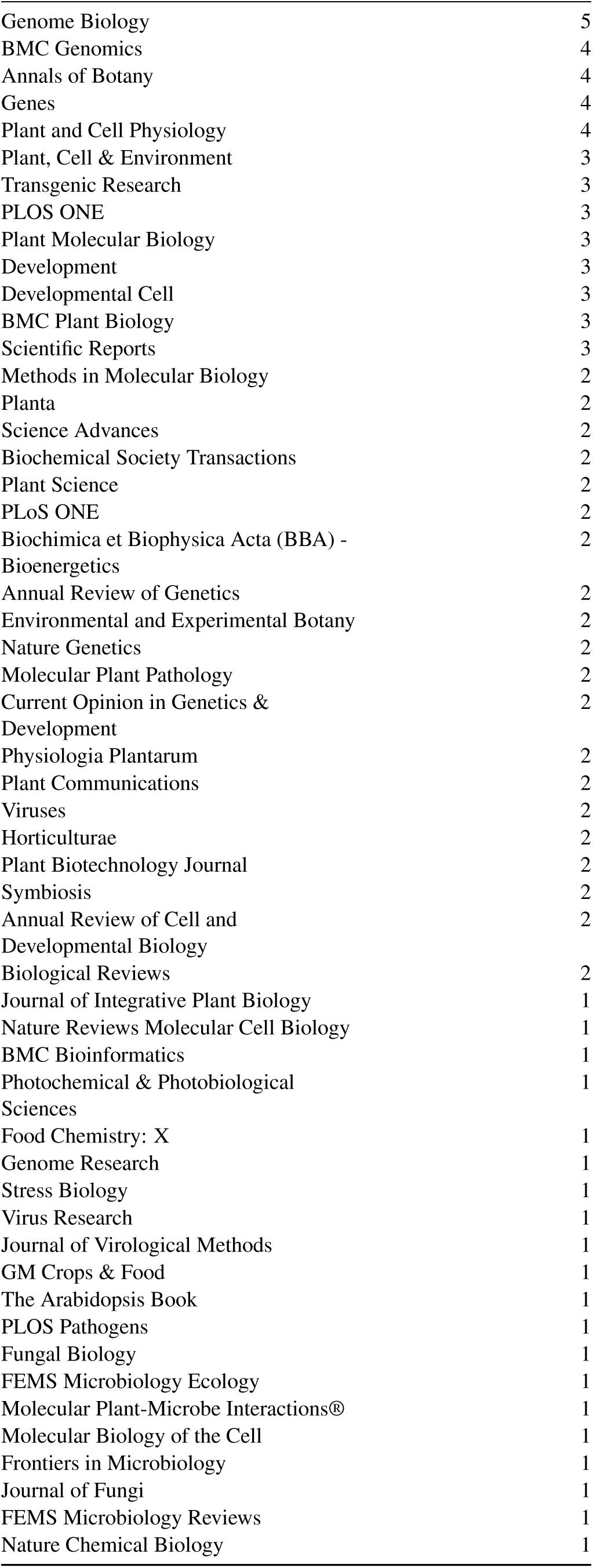
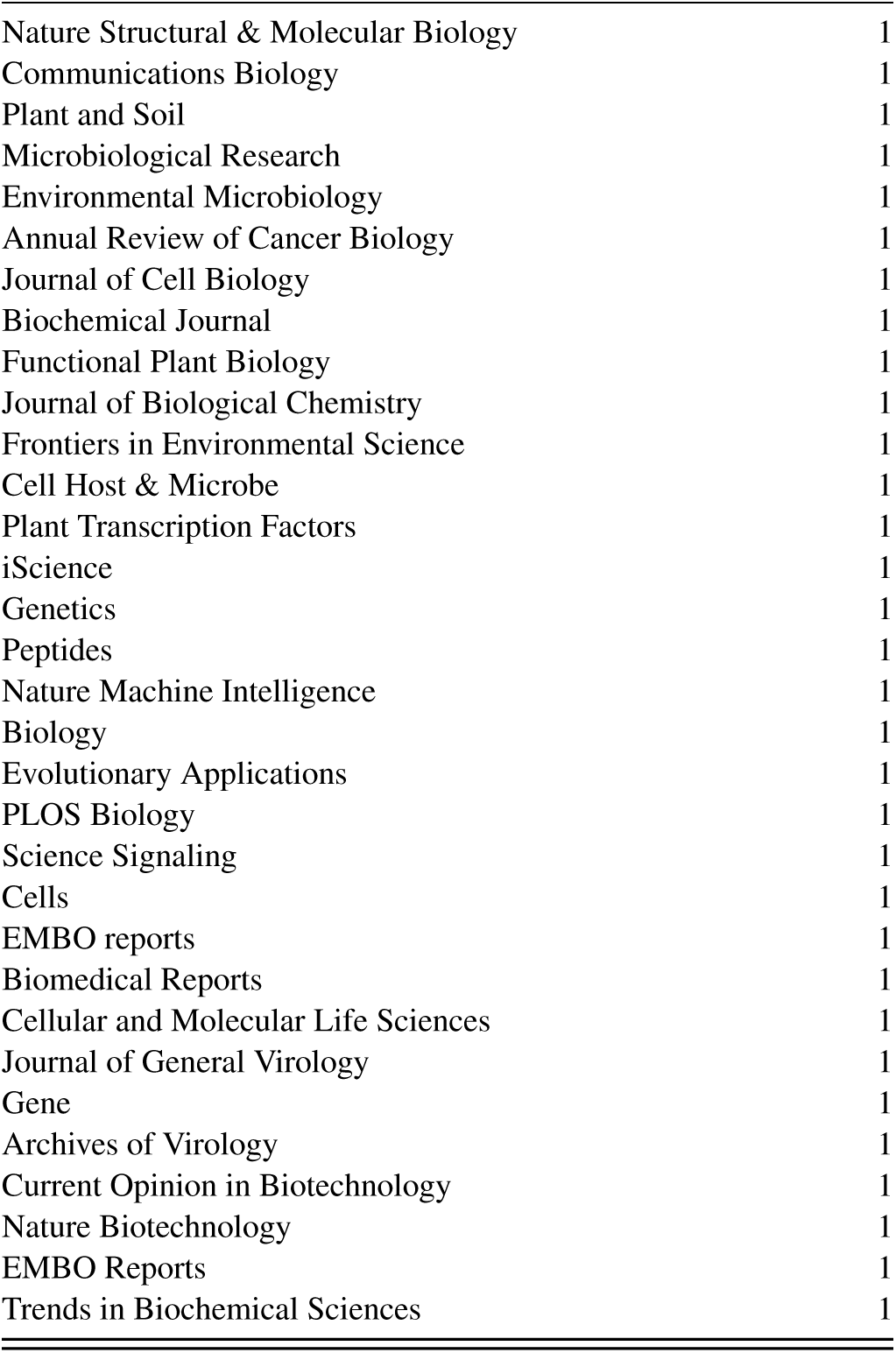
Journal distribution of the Expert MoBiPlant QA set.

| Journal | Count |
| --- | --- |
| The Plant Cell | 51 |
| Nature Communications | 35 |
| Plant Physiology | 30 |
| New Phytologist | 29 |
| The Plant Journal | 25 |
| Proceedings of the National Academy of Sciences | 24 |
| Journal of Experimental Botany | 21 |
| Nucleic Acids Research | 18 |
| Nature Plants | 16 |
| Science | 15 |
| Current Biology | 14 |
| Annual Review of Plant Biology | 13 |
| Current Opinion in Plant Biology | 12 |
| Molecular Plant | 12 |
| The EMBO Journal | 12 |
| Trends in Plant Science | 11 |
| Nature | 10 |
| Cell Reports | 10 |
| Frontiers in Plant Science | 8 |
| eLife | 8 |
| Cell | 8 |
| PLOS Genetics | 7 |
| Plants | 7 |
| International Journal of Molecular Sciences | 5 |
| Plant Cell Reports | 5 |
| Molecular Cell | 5 |

| **Journal** | **Count** |
| --- | --- |
| Genome Biology | 5 |
| BMC Genomics | 4 |
| Annals of Botany | 4 |
| Genes | 4 |
| Plant and Cell Physiology | 4 |
| Plant, Cell & Environment | 3 |
| Transgenic Research | 3 |
| PLOS ONE | 3 |
| Plant Molecular Biology | 3 |
| Development | 3 |
| Developmental Cell | 3 |
| BMC Plant Biology | 3 |
| Scientific Reports | 3 |
| Methods in Molecular Biology | 2 |
| Planta | 2 |
| Science Advances | 2 |
| Biochemical Society Transactions | 2 |
| Plant Science | 2 |
| PLoS ONE | 2 |
| Biochimica et Biophysica Acta (BBA) - Bioenergetics | 2 |
| Annual Review of Genetics | 2 |
| Environmental and Experimental Botany | 2 |
| Nature Genetics | 2 |
| Molecular Plant Pathology | 2 |
| Current Opinion in Genetics & Development | 2 |
| Physiologia Plantarum | 2 |
| Plant Communications | 2 |
| Viruses | 2 |
| Horticulturae | 2 |
| Plant Biotechnology Journal | 2 |
| Symbiosis | 2 |
| Annual Review of Cell and Developmental Biology | 2 |
| Biological Reviews | 2 |
| Journal of Integrative Plant Biology | 1 |
| Nature Reviews Molecular Cell Biology | 1 |
| BMC Bioinformatics | 1 |
| Photochemical & Photobiological Sciences | 1 |
| Food Chemistry: X | 1 |
| Genome Research | 1 |
| Stress Biology | 1 |
| Virus Research | 1 |
| Journal of Virological Methods | 1 |
| GM Crops & Food | 1 |
| The Arabidopsis Book | 1 |
| PLOS Pathogens | 1 |
| Fungal Biology | 1 |
| FEMS Microbiology Ecology | 1 |
| Molecular Plant-Microbe Interactions® | 1 |
| Molecular Biology of the Cell | 1 |
| Frontiers in Microbiology | 1 |
| Journal of Fungi | 1 |
| FEMS Microbiology Reviews | 1 |
| Nature Chemical Biology | 1 |
| Nature Structural & Molecular Biology | 1 |
| Communications Biology | 1 |
| Plant and Soil | 1 |
| Microbiological Research | 1 |
| Environmental Microbiology | 1 |
| Annual Review of Cancer Biology | 1 |
| Journal of Cell Biology | 1 |
| Biochemical Journal | 1 |
| Functional Plant Biology | 1 |
| Journal of Biological Chemistry | 1 |
| Frontiers in Environmental Science | 1 |
| Cell Host & Microbe | 1 |
| Plant Transcription Factors | 1 |
| iScience | 1 |
| Genetics | 1 |
| Peptides | 1 |
| Nature Machine Intelligence | 1 |
| Biology | 1 |
| Evolutionary Applications | 1 |
| PLOS Biology | 1 |
| Science Signaling | 1 |
| Cells | 1 |
| EMBO reports | 1 |
| Biomedical Reports | 1 |
| Cellular and Molecular Life Sciences | 1 |
| Journal of General Virology | 1 |
| Gene | 1 |
| Archives of Virology | 1 |
| Current Opinion in Biotechnology | 1 |
| Nature Biotechnology | 1 |
| EMBO Reports | 1 |
| Trends in Biochemical Sciences | 1 |

**Supplementary Table 8.** Journal distribution and model performance on the Synthetic MoBiPlant set. Answer accuracy is reported for every model.

| Journal | Count | LLaMA 3.1 405B | Claude 3.5 Sonnet | DeepSeek V3 | DeepSeek R1 | GPT-4o | o1-mini | Gemini 1.5 Pro |
| --- | --- | --- | --- | --- | --- | --- | --- | --- |
| *COPB* | 125 | 96,7 | 99,2 | 99,2 | 97,6 | 98,4 | 96,8 | 91,2 |
| *TIPS* | 135 | 96,3 | 97,8 | 97,8 | 97,8 | 97 | 96,3 | 95,6 |
| *JExBot* | 125 | 82,1 | 83,2 | 82,4 | 79,2 | 76,8 | 78,4 | 80 |
| *Molecular Plant* | 90 | 89,2 | 88,9 | 90 | 86,5 | 86,7 | 84,4 | 85,6 |
| *Nature Plants* | 55 | 75,9 | 72,7 | 80 | 75,9 | 80 | 72,7 | 80 |
| *New Phytol* | 145 | 84,8 | 76,1 | 79,6 | 78,2 | 82,4 | 82,4 | 80,3 |
| *Plant Cell* | 145 | 86,4 | 89 | 92,4 | 86,5 | 82,1 | 84,1 | 84,8 |
| *Plant Journal* | 130 | 90,4 | 89,2 | 90 | 87,3 | 86,2 | 78,5 | 81,5 |
| *Plant Phys* | 125 | 86,7 | 84,8 | 86,4 | 82,5 | 81,6 | 80,8 | 74,4 |
| **Overall** | 1075 | 87,3 | 88,4 | 89,6 | 87,3 | 85,7 | 84 | 83,9 |

## Notes

https://huggingface.co/datasets/manufernandezbur/MoBiPlant

https://github.com/manoloFer10/mobiplant

### Competing Interest Statement

The authors have declared no competing interest.

### Summary of Updates

We have added the benchmarking of GPT 5.2 and Claude Sonnet 4.6, along with a new section regarding a composite Natural Language - DNA LM (ChatNT) and tested them across both existing DNA tasks and newly crafted for this work.

